# Dissecting endothelial to haematopoietic stem cell transition by single-cell transcriptomic and functional analyses

**DOI:** 10.1101/2020.01.18.910356

**Authors:** Siyuan Hou, Zongcheng Li, Xiaona Zheng, Yun Gao, Ji Dong, Yanli Ni, Xiaobo Wang, Yunqiao Li, Xiaochen Ding, Zhilin Chang, Shuaili Li, Yuqiong Hu, Xiaoying Fan, Yu Hou, Lu Wen, Bing Liu, Fuchou Tang, Yu Lan

**Affiliations:** Key Laboratory for Regenerative Medicine of Ministry of Education, Institute of Haematology, School of Medicine, Jinan University, Guangzhou 510632, China; State Key Laboratory of Experimental Haematology, Fifth Medical Center of Chinese PLA General Hospital, Beijing 100071, China; State Key Laboratory of Proteomics, Academy of Military Medical Sciences, Academy of Military Sciences, Beijing 100071, China; Integrated Chinese and Western Medicine Postdoctoral research station, Jinan University, Guangzhou 510632, China; Beijing Advanced Innovation Center for Genomics and Biomedical Institute for Pioneering Investigation via Convergence, College of Life Sciences, Peking University, Beijing 100871, China; Ministry of Education Key Laboratory of Cell Proliferation and Differentiation, Beijing 100871, China; Peking-Tsinghua Center for Life Sciences, Peking University, Beijing 100871, China; Guangzhou Regenerative Medicine and Health-Guangdong Laboratory (GRMH-GDL), Guangzhou 510530, China

## Abstract

Haematopoietic stem cells (HSCs) in adults are believed to be born from hemogenic endothelial cells (HECs) in mid-gestational mouse embryos. Due to rare and transient nature, the HSC-competent ECs have never been stringently identified and accurately captured, let alone their genuine vasculature precursors. Here, we firstly used high-precision single-cell transcriptomics to unbiasedly examine relevant EC populations at continuous developmental stages and transcriptomically identified putative HSC-primed HECs. Combining computational prediction and in vivo functional validation, we precisely captured HSC-competent HECs by newly constructed Neurl3-EGFP reporter mouse model, and realized enrichment further by surface marker combination. Surprisingly, endothelial-haematopoietic bi-potential was rarely but reliably witnessed in culture of single HECs. Noteworthy, primitive vascular ECs experienced two-step fate choices to become HSC-primed HECs, resolving several previously observed contradictions. Taken together, comprehensive understanding of endothelial evolutions and molecular programs underlying HSC-primed HEC specification in vivo will facilitate future investigations directing HSC production in vitro.

## INTRODUCTION

The adult haematopoietic system, consisted mainly of haematopoietic stem cells (HSCs) and their multi-lineage progenies, is believed to be derived from hemogenic endothelial cells (HECs) in mid-gestational embryos ^1, 2^. It is generally accepted that while still embedded in the endothelial layer and presenting endothelial characteristics, HECs begin to express key hemogenic transcription factor Runx1 and have hemogenic potential ^3, 4^. Different from haematopoietic progenitors, HECs lack the expression of haematopoietic surface markers, such as CD41 and CD45, which mark the population capable of generating haematopoietic progenies when directly tested in colony-forming unit assays ^3, 5^. Haematopoietic stem and progenitor cells (HSPCs) are visualized to emerge from aortic endothelial cells (ECs) via a transient and dynamic process called endothelial-to-haematopoietic transition to form intra-aortic haematopoietic clusters (IAHCs) ^6–10^. Being located within IAHCs or to the deeper sub-endothelial layers, pre-HSCs serve as the important cellular intermediates between HECs and HSCs, featured by their inducible repopulating capacity and priming with haematopoietic surface markers ^11–15^. The specification of HSC-primed HECs is the initial and one of the most pivotal steps for vascular ECs to choose a HSC fate. However, as the precise identity of HSC-primed HECs is not clear, contradictory notions regarding whether primordial or arterial fated ECs are the direct origin of HSC-primed HECs are still on the debate. It is proposed that definitive HECs and arterial ECs represent distinct lineages ^16, 17^. Moreover, HSCs and arterial ECs are proposed to arise from distinct precursors, characterized by different Notch signaling strengths ^18^. Most recently, HSC-primed HECs have been transcriptionally identified in human embryos, which present an unambiguous arterial property, indicative of their arterial EC origin ^19^.

In order to deeply investigate the cellular evolutions and molecular events underlying the specification of HSC-primed HECs and their subsequent commitment to HSPCs, it is necessary to efficiently isolate the HSC-primed HECs, which is proven to be difficult not only because the population is proposed to be small and transient, but also due to the technical challenging to determine their HSC competence ^20^. Considering that not only HSCs but also the transient definitive haematopoiesis during embryogenesis are derived from HECs, repopulating capacity is required for the functional evaluation of the HSC-primed HECs. Previous studies have reported the HSC competence of CD47^+^ but not CD47^-^ ECs in embryonic day (E) 10.5 aorta-gonad-mesonephros (AGM) region and both Dll4^+^ and Dll4^-^ ECs in E9.5 para-aortic splanchnopleura (P-Sp) region ^11, 21^. Nevertheless, the enrichment of the above surface markers is far from efficient. Several transgenic reporter mouse models have been established by which HECs could be distinguished from non-HECs, including *Ly6a-GFP*, GFP transgenic reporter under the control of Runx1 +23 enhancer (*Runx1 +23GFP*) and *Gfi1-Tomato*, and the usage of these reporters largely helps to delineate the process of endothelial-to-haematopoietic transition ^3, 14, 22–25^. Although expected to a certain extent, the HSC competence of the HECs labeled by these reporters has not been functionally validated. Up-to-date, efficient isolation of the HSC-primed HEC population has not yet been achieved.

With the aim of delineating the molecular events underlying HSC emergence, several single-cell transcriptional profiling studies on HECs, IAHC cells, and HSPCs in the AGM region have been reported in recent years. Using either Runx1 +23GFP or Gfi1-Tomato as the marker of putative HECs, several defined cell populations are transcriptionally profiled by Fluidigm single-cell qPCR or single-cell RNA sequencing (scRNA-seq) ^3, 14, 22^. Moreover, the cellular components of IAHCs are investigated at single-cell level by mechanically picking up single whole IAHCs in the aortas, showing cells with pre-HSC feature are predominantly involved ^14^. Interestingly, contradiction still exists regarding whether HECs and non-HECs are molecularly similar and to what extent the two populations are distinguishable ^14, 22^. Since the enrichment efficiency or specificity of the above markers to define the HEC population might be not enough, an unsupervised screening of the embryonic endothelial pool within haematopoietic tissues is required for the precise recognition of HSC-primed HECs.

Here, we firstly used high-precision single-cell transcriptomics to unbiasedly examine all the EC populations spanning continuous developmental stages covering the presumed time points for the specification of HSC-primed HECs, transcriptomically identified them, and computationally screened for their candidate markers. Based on the consequently precise capture and isolation of the HSC-competent HECs using surface marker combination or newly constructed fluorescent reporter mice, we further decoded the cellular evolutions and molecular programs underlying the stepwise hemogenic fate settling from the initial primordial vascular ECs. A series of new findings, including the endothelial-haematopoietic bi-potential of HECs and the multi-step fate choice for the specification of HSC-primed HECs, unprecedentedly enrich our understanding of HSC generation in vivo and should be extremely critical to inspire new approaches for stepwise HSC regeneration from pluripotent stem cells ^26^.

## RESULTS

### Transcriptomic identification of the HECs in AGM region

We first analyzed mouse embryos from E9.5, when the initial IAHC formation in the aorta occurs ^7^, to the stage of the appearance of HSCs at E11.0 ^27^ (Supplementary information, Fig. S1a). For each embryo, the embryo proper was isolated and the head, limb buds, heart, visceral bud, and vitelline and umbilical vessels outside the embryo proper were excluded (Fig. 1a). To specifically capture aortic luminal ECs of AGM region, we performed microinjection of fluorescent dye Oregon green into the dorsal aortas of E10.0-E11.0 embryos as reported ^12^ (Fig. 1a; Supplementary information, Fig. S1b). The sampled cells were purified by FACS as CD45^-^CD31^+^CD144^+^, which contained predominantly vascular ECs and CD41^+^ haematopoietic cells. Meanwhile, CD45^-^CD31^-^CD144^-^ non-EC cells in the body were used as negative controls (Fig. 1a). We used unique molecular identifier (UMI)-based scRNA-seq method to accurately measure the gene expression profiles within individual cells. In totally 662 sequenced single cells, 597 single-cell transcriptomes passed rigorous quality control. On average we detected 7,035 genes (from 2,266 to 10,843) and 636,418 transcripts (from 103,793 to 2,959,573) expressed in each individual cell (Supplementary information, Fig. S1c).

**Fig. 1.**
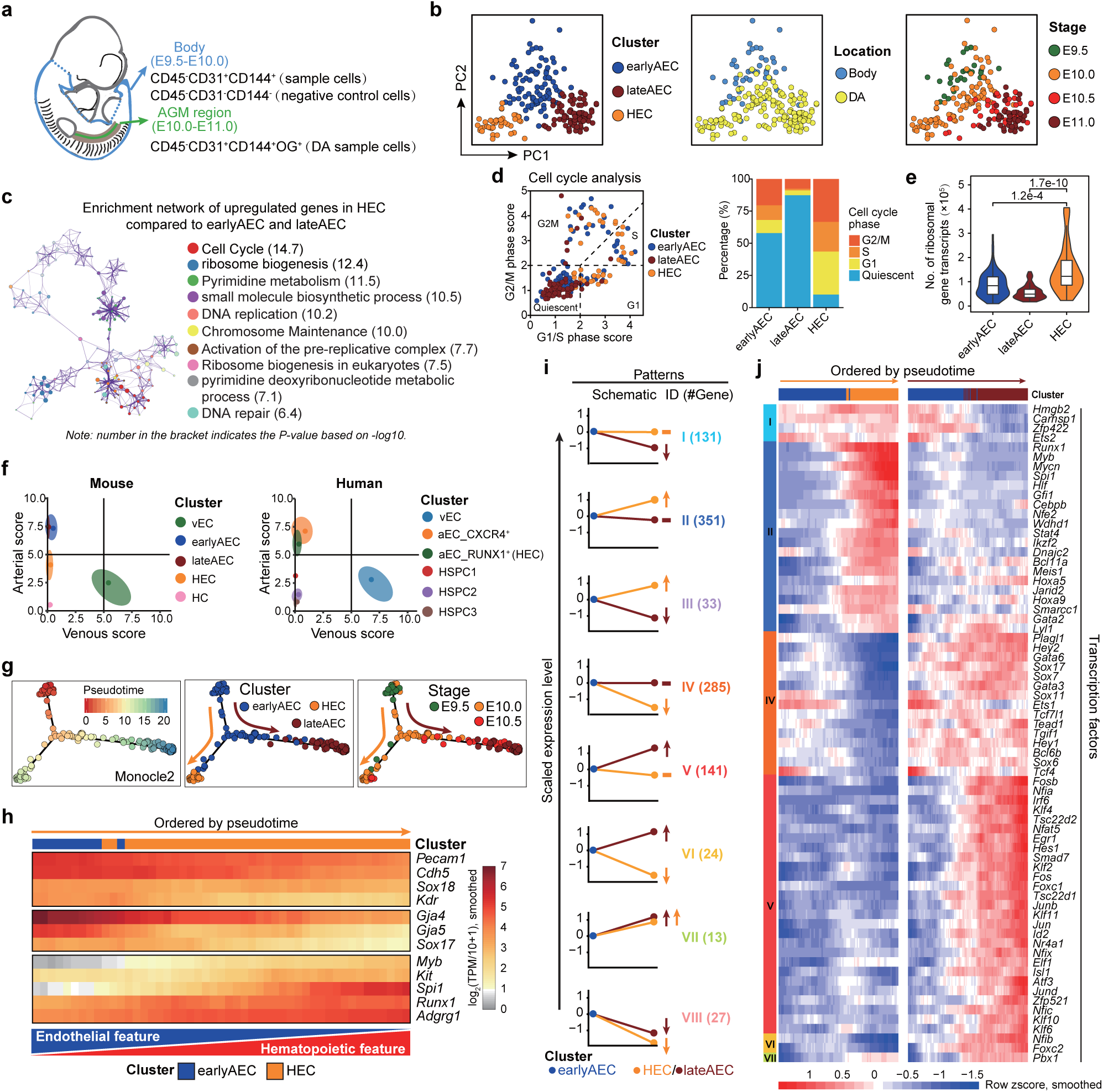
Transcriptomic identification and molecular characteristics of the HECs in AGM Region. (a) Schematic illustration of the strategies used for embryo dissection and cell preparation for the subsequent scRNA-seq. The involved body part and AGM region is indicated as blue and green, respectively, with head, limb buds, heart, visceral bud, and umbilical and vitelline vessels outside the embryo proper excluded. (b) PCA plots with three clusters (earlyAEC, lateAEC and HEC) (left), sampling locations (middle) and embryonic stages (right) mapped onto it. (c) Metascape network enrichment analysis with top 10 enriched terms exhibited to the right. Each cluster is represented by different colors and each enriched term is represented by a circle node. Number in the bracket indicates the *P* value based on -log10. (d) Classification of the indicated cells into quiescent phase and other cycling phases (G1, S and G2M) based on the average expression of G1/S and G2/M gene sets (left). Stacked bar chart showing the constitution of different cell cycle phases in the corresponding three clusters shown to the left (right). (e) Violin plot showing the number of transcripts for ribosomal related genes detected in each single cell of the indicated clusters. Wilcoxon Rank Sum test is employed to test the significance of difference and *P* values are indicated for the comparison. *P* < 0.05 is considered statistically significant. (f) Scatterplot showing the average arteriovenous scores of the cells in each cluster for mouse dataset in this paper (left) and human dataset from published articles (right), respectively. Main distribution ranges of arteriovenous scores in each cluster are also indicated as an oval shape. (g) Pseudotemporal ordering of the cells in select three clusters inferred by monocle 2, with pseudotime (left), clusters (middle) and sampling stages (right) mapped to it. HEC specification and AEC maturation directions are indicated as orange and deep red arrows, respectively. (h) Heatmap showing the expression of the indicated genes (smoothed over 15 adjacent cells) with cells ordered along the pseudotime axis of HEC specification branch inferred by monocle 2. (i) Eight major expression patterns identified from the differentially expressed genes in HEC or lateAEC as compared to earlyAEC. Arrows showing the changes in HEC or lateAEC as compared to earlyAEC. The numbers of pattern genes are indicated to the right. (j) Heatmaps showing the relative expressions (smoothed over 20 adjacent cells) of the TFs belonging to the pattern genes with cells ordered along the pseudotime axis and genes ordered by patterns.

According to a graph-based clustering approach from Seurat software ^28^, all cells were separated into six clusters, including one negative (Neg) cluster containing mainly non-EC negative control cells, and five sample clusters comprising almost all FACS-isolated sample cells (Supplementary information, Fig. S1d and Table S1). Featured by the obvious *Runx1* and *Itga2b* (encoding CD41) expression, the haematopoietic cell (HC) cluster was distributed away from the other four vascular EC clusters which presented apparent arterial or venous characteristics (Supplementary information, Fig. S1d, e). One venous EC (vEC) cluster was readily recognized by the exclusive expression of *Nr2f2* in all vascular EC populations (Supplementary information, Fig. S1d, e). Two arterial EC clusters showed similar *Gja5* expression but different level of *Ltbp4* expression ^29^. Together with their different sampling stages (mainly from E9.5-E10.0 and E10.5-E11.0, respectively), they were annotated as early arterial EC (earlyAEC) and late arterial EC (lateAEC) cluster, respectively (Supplementary information, Fig. S1d, e). The left one cluster basically met the criteria of the molecular definition of HEC, showing apparent *Runx1* expression upon endothelial property, and was consequently named as HEC cluster (Supplementary information, Fig. S1d, e). To more strictly define the HEC population, cells within Neg cluster and those transcriptionally expressing *Ptprc* (encoding CD45) or *Spn* (encoding CD43) were excluded for the subsequent analysis (Supplementary information, Fig. S1f).

HEC and other two AEC clusters were further focused as they were either molecularly or spatiotemporally near to each other (Fig. 1b, Supplementary information, Fig. S1d). To exclude the possibility that we failed to identify important populations relevant to hemogenic specification in earlyAEC cluster, which contributed evidently to the aortic inner layer of AGM region at E10.0 (Supplementary information, Fig. S1f), we performed forced clustering within the given cluster. *Runx1* (signature of hemogenic specification) was not significantly differentially expressed between the two sub-clusters, suggesting that no population with sign of hemogenic specification was missed by our clustering (Supplementary information, Fig. S1g). Moreover, very few genes were significantly differentially expressed in the forced sub-clusters of HEC, and none of them was related to hemogenic or haematopoietic features, indicative of the largely homogeneous property of the HEC cluster (Supplementary information, Fig. S1g). HEC was reduced promptly in number at E10.5, and became hardly detectable by E11.0 (Fig. 1b; Supplementary information, Fig. S1f). The highly expressed genes in HEC as compared to earlyAEC and lateAEC were mainly enriched in the terms related to cell cycle and ribosome biogenesis (Fig. 1c; Supplementary information, Table S2). Cell cycle analysis demonstrated a remarkably activated cycling in HEC, in sharp contrast to the quiescent state by arterial EC maturation (Fig. 1d). On average, each cell in HEC cluster expressed more mRNA molecules and ribosomal genes than either earlyAEC or lateAEC (Fig. 1e; Supplementary information, Fig. S1h), supportive of the globally up-regulated transcriptional and translational activity during hemogenic specification, which was in line with the finding in human embryo that translational initiation is overrepresented in HSC-primed HECs than in arterial ECs ^19^. We further evaluated the arteriovenous scores of the populations we defined, and found similar results in mouse and human that HEC rather than haematopoietic populations manifested certain arterial feature (Fig. 1f).

Trajectory analysis by Monocle 2 suggested that along the arterial maturation path from earlyAEC towards lateAEC, HEC was segregated out from earlyAEC at E9.5-E10.0 (Fig. 1g). The gradual up-regulation of hemogenic genes, including *Runx1* and *Spi1*, was accompanied by the gradual down-regulation of both endothelial and arterial genes along the HEC specification pseudotime, with the endothelial-haematopoietic dual-feature of the HEC population presenting as a dynamic continuum (Fig. 1h). The finding was in line with previous report about the reciprocal expression of Runx1 and Sox17 in HECs ^30^. To search for the genes that would be potentially meaningful to the distinct fate choices of earlyAEC, those differentially expressed between earlyAEC and its downstream population HEC or lateAEC were screened out, and eight major patterns were witnessed (Fig. 1i; Supplementary information, Fig. S1i and Table S3). Most of these genes showed altered expression along one but not both specification paths from earlyAEC (Pattern I, II IV, and V) (Fig. 1i; Supplementary information, Fig. S1i). Most transcription factors (TFs) within these patterns were those up-regulated along either HEC specification or arterial EC maturation (Fig. 1j; Supplementary information, Fig. S1j). Interestingly, both *Hoxa5* and *Hoxa9* belonged to the same pattern as *Runx1*, although their expression was not well-correlated with *Runx1* (Fig. 1j; Supplementary information, Fig. S1j, k). The data suggested that the gene expressions should be orchestrated and precisely regulated for the subsequent cell fate choice from earlyAEC.

### Efficient capture of the HSC-competent and endothelial-haematopoietic bi-potent HECs in AGM region

We next made an effort to identify surface marker combination to highly enrich the HECs for functional evaluation (Supplementary information, Table S4). *Cd44*, *Procr* (coding CD201) and *Kit* were screened out by differentially expressed genes and correlation analysis (Fig. 2a). We specifically focused on E10.0 in the following functional assays to keep consistent with the transcriptomic finding. Whole-mount immunostaining showed that in addition to the scattered blood cells throughout the tissue, the expression of CD44 was detected in the whole endothelial layer of dorsal aorta and very proximal part of its segmental branches (Fig. 2b). Using similar strategy as for pre-HSC identification ^11^, we found only the derivatives induced from CD41^-^CD43^-^CD45^-^CD31^+^Kit^+^CD201^+^ rather than CD41^-^CD43^-^CD45^-^CD31^+^Kit^+^CD201^-^ population at E9.5-E10.0 could long-term (16 weeks) and multi-lineage reconstitute lethally irradiated adult recipients, although both populations generated haematopoietic clusters with different frequencies upon 7 days culture on OP9-DL1 stromal cells (Fig. 2c-e; Supplementary information, Fig. S2a-c). Self-renewal capacity of the HSCs was further validated by secondary transplantation (Fig. 2d, e; Supplementary information, Fig. S2b). Within CD41^-^CD43^-^CD45^-^CD201^+^ population, induced HSC potential was exclusively detected in CD44^+^ subpopulation (Fig. 2c-e; Supplementary information, Fig. S2a-c). Thus, our data identified CD41^-^CD43^-^CD45^-^CD31^+^CD201^+^Kit^+^CD44^+^ (PK44) population in E10.0 caudal half as the HSC-competent HECs.

**Fig. 2.**
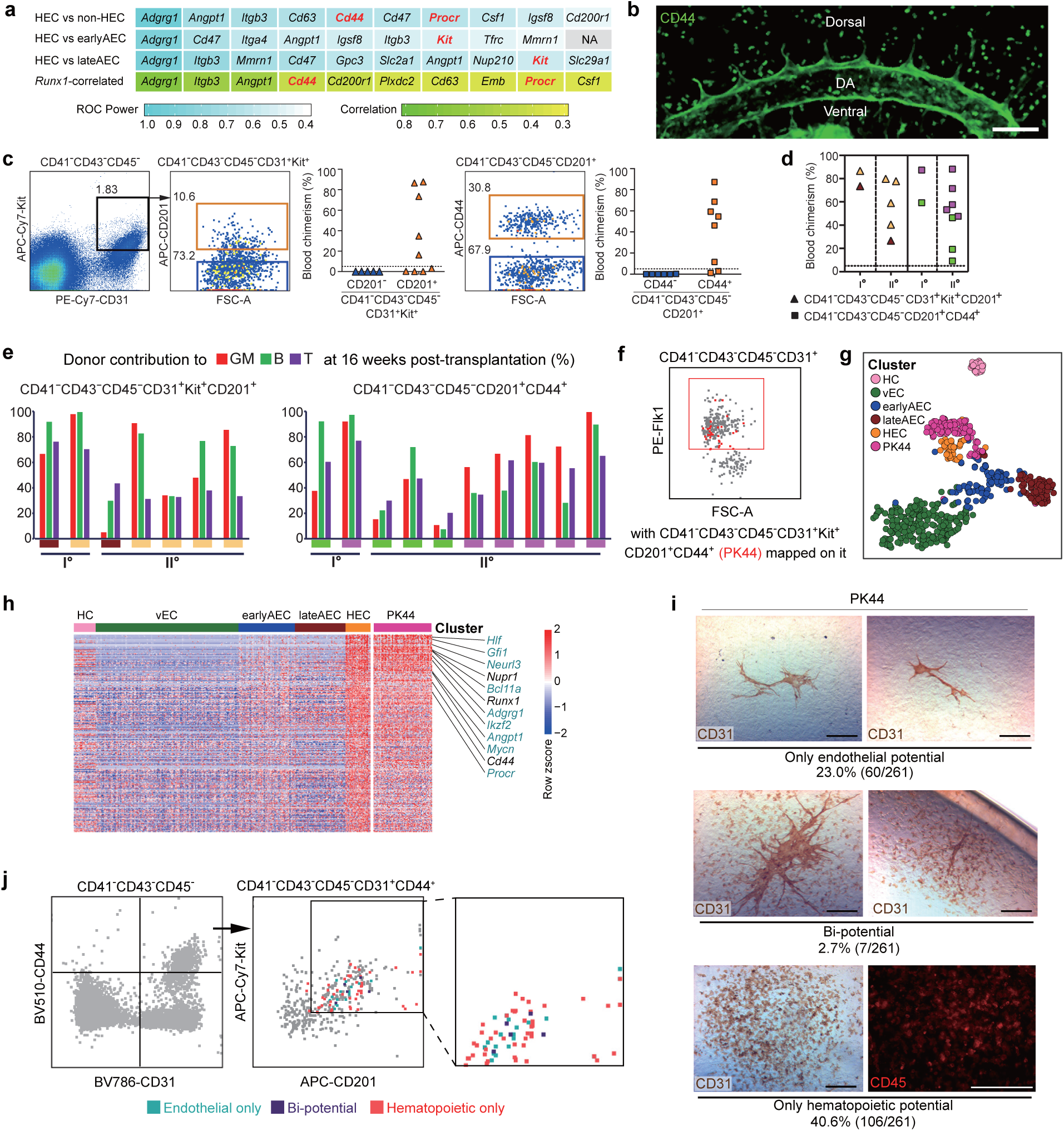
Efficiently isolating the HSC-competent and endothelial-haematopoietic bi-potent HECs before HSC emergence. (a) Gene lists of the top ten cell surface molecules significantly overrepresented in HEC as compared to the indicated cell populations (first 3 lines) and those positively correlated with Runx1 within 4 EC clusters (vEC, earlyAEC, lateAEC and HEC, last line). Non-HEC, cells except for HEC within 4 EC clusters. Highlights in red font indicate the candidates used for further functional analysis. (b) Representative whole-mount staining of CD44 at E10.0 AGM region, showing CD44 is expressed in the whole endothelial layer of the dorsal aorta and roots of its proximal branches. DA, dorsal aorta; Scale bar, 100 μm. (c) Representative FACS plots for cell sorting of the E9.5-E10.0 caudal half for co-culture/transplantation assay and the donor chimerism at 16 weeks after transplantation of the derivatives of the indicated cell populations. (d) Blood chimerism of the primary (I°) and corresponding secondary (II°) recipients at 16 weeks post-transplantation. The primary recipients were transplanted with the derivatives of the indicated cells from the caudal half of E9.5-E10.0 embryos. The paired primary and corresponding secondary repopulated mice are shown as the same symbol and color. (e) Bars represent the percent donor contribution to the granulocytes/monocytes (GM, red), B lymphocytes (green), and T lymphocytes (purple) in the peripheral blood of the primary (I°) and secondary (II°) recipients at 16 weeks post-transplantation. The paired primary and corresponding secondary repopulated mice are shown as the same colors below. (f) FACS plot of Flk1 expression in the indicated population of E10.0 AGM region, with PK44 (CD41^-^CD43^-^CD45^-^CD31^+^CD201^+^Kit^+^CD44^+^) cells (red) mapped onto it. Red box indicates the gate of Flk1^+^ cells. (g) t-SNE plot of the cells included in the filtered initial dataset and PK44 dataset, with clusters mapped on it. PK44, CD41^-^CD43^-^CD45^-^CD31^+^CD201^+^Kit^+^CD44^+^ population from E10.0 AGM region. (h) Heatmap showing the relative expressions of HEC feature genes, which are defined as those significantly highly expressed as compared to others including HC, vEC, earlyAEC and lateAEC, in the indicated cell populations. Selected HEC feature genes are shown to the right with pre-HSC signature genes marked as aquamarine. (i) Representative CD31 and CD45 immunostaining on the cultures of single PK44 cells from E10.0 AGM region, showing typical morphologies regarding distinct differentiation potentials. Cell frequencies of each kind of potential are also shown. Data are from 5 independent experiments with totally 15 embryos used. Scale bars, 400 μm. (j) Expression of Kit and CD201 in the index-sorted single PK44 cells with differentiation potential based on in vitro functional evaluation. Cells with different kinds of potential are mapped onto the reference FACS plots (grey dots). Box in the middle plot indicates the gate for FACS sorting of PK44 cells in E10.0 AGM region and its enlarged view is shown to the right.

As compared to other endothelial surface markers, Flk1 (encoded by *Kdr*) is known to be specifically localized within the vessel lumen layer, and very few and only the basal-most localized IAHC cells express Flk1 ^7^. Here, almost all PK44 cells expressed Flk1 by FACS analysis, indicative of their endothelial layer localization (Fig. 2f). To determine the transcriptomic identity of the HSC-competent HECs we isolated, totally 96 PK44 single cells derived from E10.0 AGM were sequenced (Supplementary information, Table S1). The PK44 cells were clustered together with HEC by computational assignment (Fig. 2g), and showed a similar high expression of the HEC feature genes (Fig. 2h). The ubiquitous and obvious expression of several key haematopoietic TFs, including *Runx1*, *Spi1*, *Gfi1*, and *Myb* ^3, 22^, in PK44 cells inferred the enrichment of hemogenic potential (Fig. 2h; Supplementary information, Fig. S2d). Therefore, immunophenotypically purified PK44 cells elegantly represented the transcriptomically defined HEC.

We next explored whether endothelial-haematopoietic bi-potential existed in these HSC-competent HECs, since that if a cell population is experiencing fate choice so that the transient intermediate state might be captured. Firstly, we found that the Kit^+^CD201^+^ ECs in the body part of embryo proper at E9.5-E10.0 had a relatively higher endothelial tube-forming capacity as compared to the Kit^-^ or Kit^+^CD201^-^ endothelial populations (Supplementary information, Fig. S2e). Furthermore, CD44^+^ and CD44^-^ fractions within Kit^+^CD201^+^ ECs showed comparable endothelial tube-forming capacity whereas the generation of haematopoietic cells in the cultures was exclusively detected in the CD44^+^ ones under the bi-potential induction system (Supplementary information, Fig. S2f). The data suggested the largely maintenance of endothelial potential in the HECs. By single-cell in vitro induction, 40.6% (106/261) PK44 cells gave rise to only haematopoietic progenies and 23.0% (60/261) only endothelial tubules. Remarkably, 2.7% (7/261) had both haematopoietic and endothelial potential (Fig. 2i; Supplementary information, Fig. S2f). All three kinds of potential did not present an obviously biased distribution regarding Kit or CD201 expression level by index sorting analysis (Fig. 2j). Such rare bi-potential should properly represent the intermediate cellular state in HECs along their specification path (Fig. 1h), with both endothelial and haematopoietic competencies being reflected by the asymmetric cell division under in vitro culture condition, which further emphasized the efficiency of capturing such a dynamic functional population via unsupervised computational screening.

### Transcriptional and functional relationship between HECs and T1 pre-HSCs

Since the transcriptomically identified HECs and the immunophenotypically defined HSC-competent HECs (PK44) presented a largely similar molecular features (Fig. 2g, h; Supplementary information, Fig. S2d), we combined them as transcriptomic & immunophenotypic & functional HEC (tif-HEC) for the subsequent analysis. tif-HEC expressed a series of pre-HSC signature genes we previously identified ^11^, including *Hlf*, *Gfi1*, *Neurl3*, *Bcl11a*, *Adgrg1*, *Ikzf2*, *Angpt1*, *Mycn*, and *Procr*, suggestive of their HSC-related identity (Fig. 2h). We further performed scRNA-seq of 47 T1 pre-HSCs (CD31^+^CD45^-^CD41^low^Kit^+^CD201^high^) from E11.0 AGM using the same sequencing strategy as other cells in the present study ^11^ (Fig. 3a; Supplementary information, Table S1). As compared to tif-HEC, T1 pre-HSC expressed similar level of *Runx1* and *Gfi1* but obviously higher level of *Spn* (encoding CD43), validating its haematopoietic cell identity (Fig. 3b). The distribution of most T1 pre-HSCs was adjacent to tif-HEC via t-SNE visualization (Fig. 3c). Of note, principal component (PC) 2 by PCA analysis largely captured the transcriptomic differences between tif-HEC and T1 pre-HSC (Fig. 3d). The genes enriched in PC2 positive direction, where tif-HECs were mainly localized, were related to cell division, vascular development and cell spreading (Fig. 3e). Consistently, approximately 90% cells in HEC were proliferative (Fig. 1d), whereas the constitution is only about half in the T1 pre-HSCs ^11^. Serving as the extracellular matrix component of blood vessels, *Col4a1* was expressed higher in tif-HEC than in T1 pre-HSC, further confirming the vascular endothelial property of the HECs we identified ^31^ (Fig. 3f). In comparison, the genes enriched in PC2 negative direction mainly related to RNA splicing and blood coagulation (Fig. 3e). Together with the overrepresented *Spi1* in T1 pre-HSC (Fig. 3f), the data suggested that haematopoietic activity has been activated in T1 pre-HSC as compared to HEC.

**Fig. 3.**
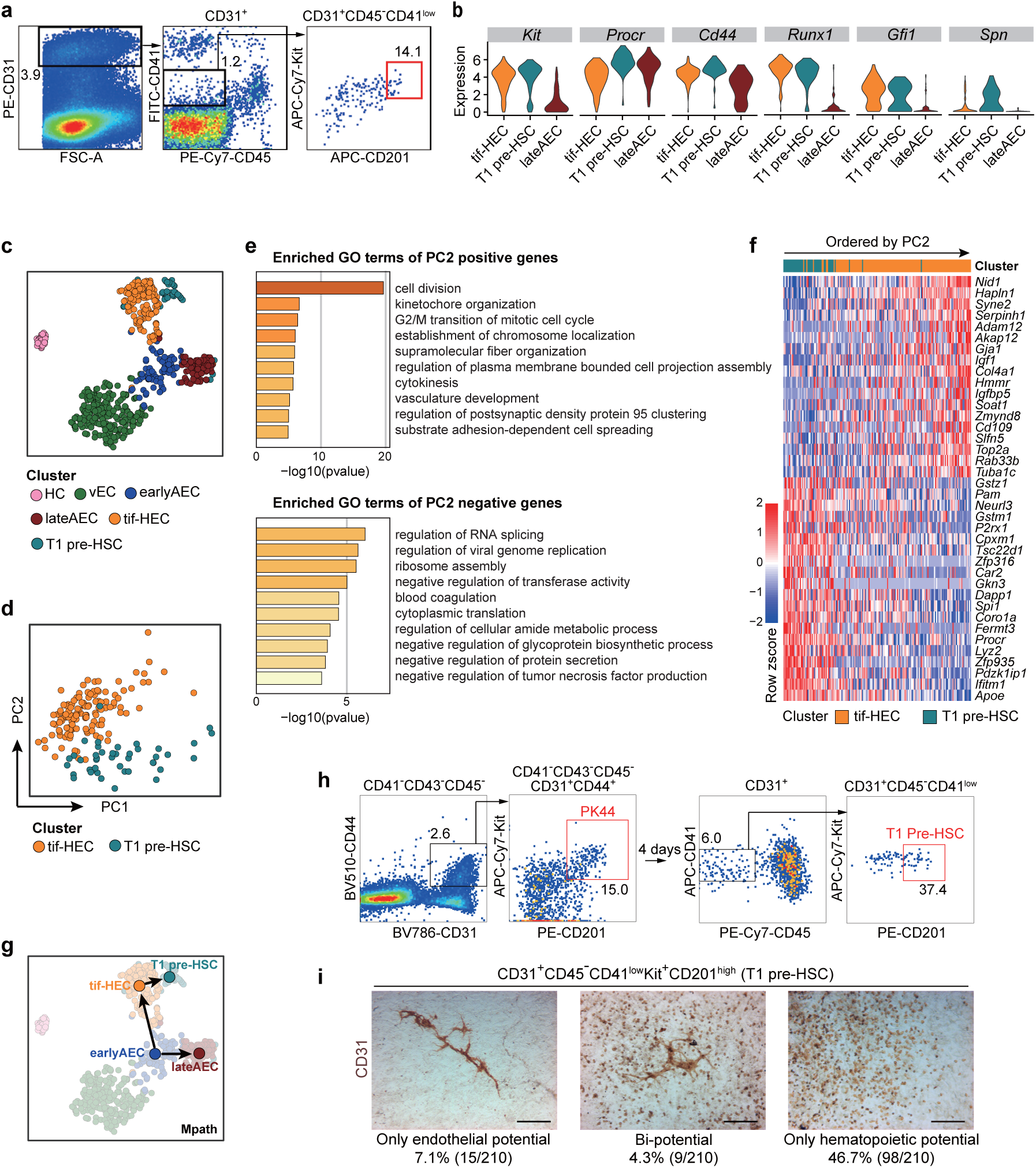
Relationship between HSC-primed HECs and T1 pre-HSCs. (a) Representative FACS plots for sorting of the T1 pre-HSCs (CD31^+^CD45^-^CD41^low^Kit^+^CD201^high^) from E11.0 AGM region of mouse embryos. Red box indicates the sampling cells for scRNA-seq. (b) Violin plots showing the expression levels of indicated genes in tif-HEC (including cluster HEC and PK44), T1 pre-HSC and lateAEC. (c) t-SNE plot of the cells included in the filtered initial dataset, PK44 dataset and T1 pre-HSC dataset, with clusters mapped on it. Cluster HEC and PK44 are combined as tif-HEC. (d) PCA plot of tif-HEC and T1 pre-HSC populations. (e) Enriched terms of PC2 positive and negative genes are shown, corresponding to the properties distinguishing tif-HEC and T1 pre-HSC, respectively. (f) Heatmap showing top 20 positive and negative genes of PC2. Genes were ordered by their contributions to PC2. (g) Trajectory of AEC clusters, tif-HEC and T1 pre-HSC inferred by Mpath. Arrows indicate the development directions predicted by sampling stages. (h) Representative FACS plots for sorting of the PK44 cells from E10.0 AGM region (left) and analysis of the immunophenotypic T1 pre-HSCs (right) after cultured in vitro for 4 days. (i) Representative CD31 immunostaining on the cultures of single T1 pre-HSCs from E11.0 AGM region, showing typical morphologies regarding distinct differentiation capacities. Cell frequencies of each type are also shown. Data are from 7 independent experiments with totally 89 embryos used. Scale bars, 400 μm.

The developmental path from tif-HEC to T1 pre-HSC was inferred by Mpath trajectory analysis (Fig. 3g). Consistently, during the course of in vitro culture of the PK44 population from E10.0 AGM region on OP9-DL1 stromal cells to induce its HSC activity, we could witness the generation of immunophenotypic T1 pre-HSCs (Fig. 3h). We also evaluated the endothelial and haematopoietic potentials of the T1 pre-HSCs (CD31^+^CD45^-^CD41^low^Kit^+^CD201^high^) in E11.0 AGM region at single-cell level. Surprisingly, we found that although displayed largely decreased endothelial potential as compared to E10.0 PK44 cells (Fig. 2i), T1 pre-HSCs still maintained comparable endothelial-haematopoietic bi-potential as that in PK44 population (Fig. 3i). This finding implied that the extremely rare and enriched T1 pre-HSC population has not completely fulfilled the endothelial-to-haematopoietic fate transition.

### Enrichment of the HSC-competent HECs by newly established Neurl3-EGFP reporter

In an effort to search for single markers to distinguish HSC-primed HECs from non-HECs or those CD45^-^CD43^-^ haematopoietic cells sharing an endothelial immunophenotype, we computationally screened for the genes significantly overrepresented in HEC cluster as compared to each of the other four clusters, including one haematopoietic cluster (HC) and three vascular EC clusters (vEC, earlyAEC and lateAEC) (Supplementary information, Fig. S1f). Totally eleven genes were screened out, which were then designated as signature genes of HSC-primed HEC, including three TFs (*Mycn*, *Hlf* and *Gfi1*) but no cell surface markers (Fig. 4a; Supplementary information, Table S5). Most of them manifested similarly high expression in T1 pre-HSCs, with six of them, namely *Neurl3*, *Dnmt3b*, *Mycn*, *Hlf*, *Gfi1*, and *Gck*, belonged to pre-HSC signature genes ^11^ (Fig. 4a).

**Fig. 4.**
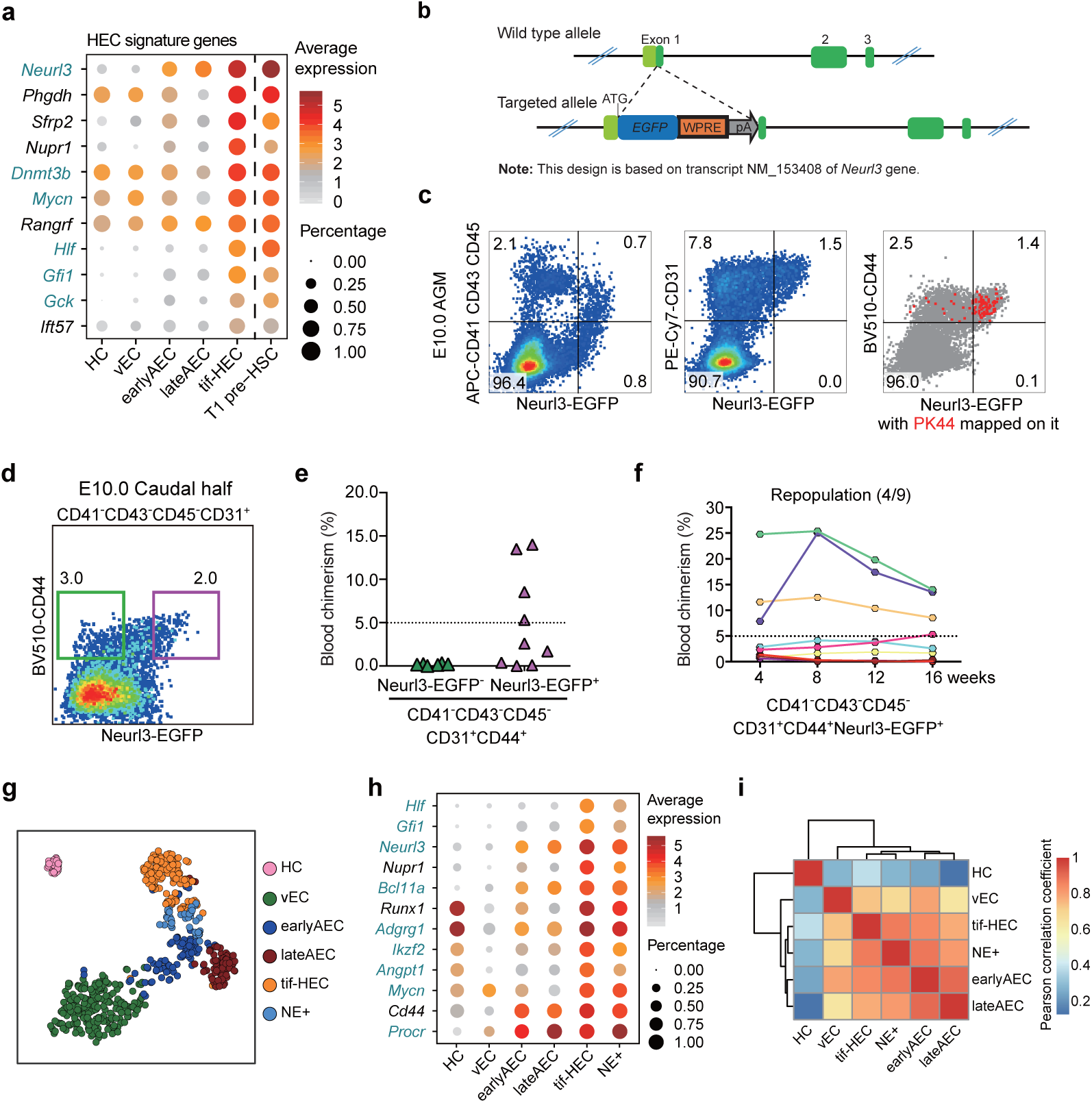
Identifying Neurl3 as a signature gene of HSC-primed HECs validated by functional and transcriptomic evaluation. (a) Dot plot showing the average and percentage expression of HEC signature genes in the indicated clusters. Genes are ordered by their median expression level in tif-HEC. Pre-HSC signature genes are marked as aquamarine. (b) Schematic model of the gene targeting strategy for generating *Neurl3^EGFP/+^* reporter mouse line via CRISPR/Cas9 system. (c) Representative FACS analysis of the E10.0 AGM region in *Neurl3^EGFP/+^* embryos, FACS plot to the right showing PK44 cells (red dots) mapped on it. (d) Representative FACS plot for sorting of the indicated cell populations from E10.0 caudal half of *Neurl3^EGFP/+^* embryos. (e) Graph showing the donor chimerism at 16 weeks after transplantation of the derivatives of the indicated populations from the caudal half of E10.0 *Neurl3^EGFP/+^* embryos. (f) Graph showing the donor chimerism at 4-16 weeks post-transplantation. The recipients were transplanted with the derivatives of CD41^-^CD43^-^CD45^-^CD31^+^CD44^+^Neurl3-EGFP^+^ population from the caudal half of E10.0 *Neurl3^EGFP/+^* embryos. Number of repopulated/total recipients is shown in the brackets. (g) t-SNE plot of the cells included in the filtered initial dataset and additional PK44 and NE+ datasets, with clusters mapped on it. Cluster HEC and PK44 are combined as tif-HEC. NE+, CD41^-^CD43^-^CD45^-^CD31^+^CD44^+^Neurl3-EGFP^+^ population from E10.0 AGM region. (h) Dot plot showing the average and percentage expression of selected HEC feature genes in the indicated clusters. Pre-HSC signature genes are marked as aquamarine. (i) Heatmap showing the correlation coefficient between each two clusters with hierarchical clustering using average method. Pearson correlation coefficient is calculated using average expression of highly variable genes in each cluster.

To further validate the bioinformatics findings and precisely determine the localization of these HSC-primed HECs, we specially chose *Neurl3* to establish a fluorescence reporter mouse line with the mind of possessing enough sensitivity, as the median expression of *Neurl3* was the highest among these signature genes (Fig. 4a; Supplementary information, Table S5). By CRISPR/Cas9-mediated gene knockin strategy, the *EGFP* was inserted into the translational initiation codon of mouse *Neurl3* gene to ensure that EGFP would be expressed in exactly the same way as Neurl3 (Fig. 4b). We firstly evaluated the Neurl3-EGFP expression by flow cytometric analysis (Fig. 4c). At E10.0 AGM region, about half of the Neurl3-EGFP^+^ cells were haematopoietic (CD41/CD43/CD45-positive) cells, which constituted about one fourth of haematopoietic population (Fig. 4c). All the Neurl3-EGFP^+^ cells were CD31^+^, and nearly all of them expressed CD44, indicative of the predominant aortic localization of Neurl3-EGFP^+^ ECs (Fig. 2b, 4c). Importantly, most PK44 cells were Neurl3-EGFP^+^, highly suggesting the enrichment of HSC-competence by the Neurl3-EGFP^+^ ECs (Fig. 4c). To confirm the HSC-competence of the Neurl3-EGFP-labeled ECs, we performed co-culture plus transplantation assay using E10.0 Neurl3-EGFP mouse embryos (Fig. 4d). Although both could generate haematopoietic clusters under the in vitro cultures, all the long-term (16 weeks) repopulations were detected exclusively in the recipients transplanted with the derivatives from CD44^+^Neurl3-EGFP^+^ ECs but not from CD44^+^Neurl3-EGFP^-^ ECs (Fig. 4e, f; Supplementary information, Fig. S3a).

We next investigated the transcriptomic identity of the Neurl3-EGFP^+^ ECs and totally 48 ECs with an immunophenotype of CD41^-^CD43^-^CD45^-^CD31^+^CD44^+^Neurl3-EGFP^+^ (NE^+^) from E10.0 AGM region were sequenced. All the NE^+^ cells ubiquitously expressed EGFP as expected and most of them expressed *Nerul3* and *Runx1* (Supplementary information, Fig. S3b, c). They distributed close to tif-HEC and were predominantly located between tif-HEC and earlyAEC by t-SNE visualization (Fig. 4g; Supplementary information, Table S1). Accordingly, NE^+^ cells demonstrated the increased cycling as compared to earlyAEC, presenting an intermediate proliferative status between earlyAEC and tif-HEC (Supplementary information, Fig. S3d). Similar to tif-HEC, NE^+^ cells showed relatively high expression of a set of HEC feature genes and pre-HSC signature genes (Fig. 2h, 4h). Correlation analysis revealed that NE^+^ cells showed the highest similarity with tif-HEC and they were clustered together by hierarchical clustering, whereas earlyAEC and lateAEC were much correlated (Fig. 4i). Therefore, from the immunophenotypic, functional and transcriptomic evaluation, the performance of the Neurl3-EGFP-marked ECs was consistent with the prediction of HSC-primed HEC by unsupervised computational screening.

### In situ localization and in vitro function of the HECs marked by Neurl3-EGFP reporter

At the AGM region of E9.5-E11.0 embryos, CD44 expression marked the whole endothelial layer of dorsal aorta in addition to IAHC cells, in line with the whole mount staining (Fig. 2b, 5a; Supplementary information, Fig. S3e). Of note, Neurl3-EGFP expression was restricted to the IAHCs and partial aortic ECs, where Neurl3-EGFP and Runx1 presented a highly co-expressed pattern (Fig. 5a; Supplementary information, Fig. S3e). Thus the Neurl3-EGFP^+^ cells embedded in the endothelial layer largely enriched the putative HECs. By FACS analysis, the average constitution of Neurl3-EGFP^+^ cells in CD44^+^ ECs were 37.7%, 50.2% and 18.3% in E9.5 caudal half, E10.0 and E10.5 AGM region, respectively. Considering the slightly over-estimation due to the much sensitivity of FACS, the data were basically in accordance with the morphological finding (Fig. 5a) and the estimated HEC composition by scRNA-seq (Supplementary information, Fig. S1f). The temporal dynamics of the HEC we defined here was in line with that of Runx1 expression in aortic endothelial layer ^32^, and the peaking of which at E10.0 was about 0.5 days earlier than the time point that the number of IAHC cells reaches the peak and the first HSCs are detected in AGM region ^7, 27^.

**Fig. 5.**
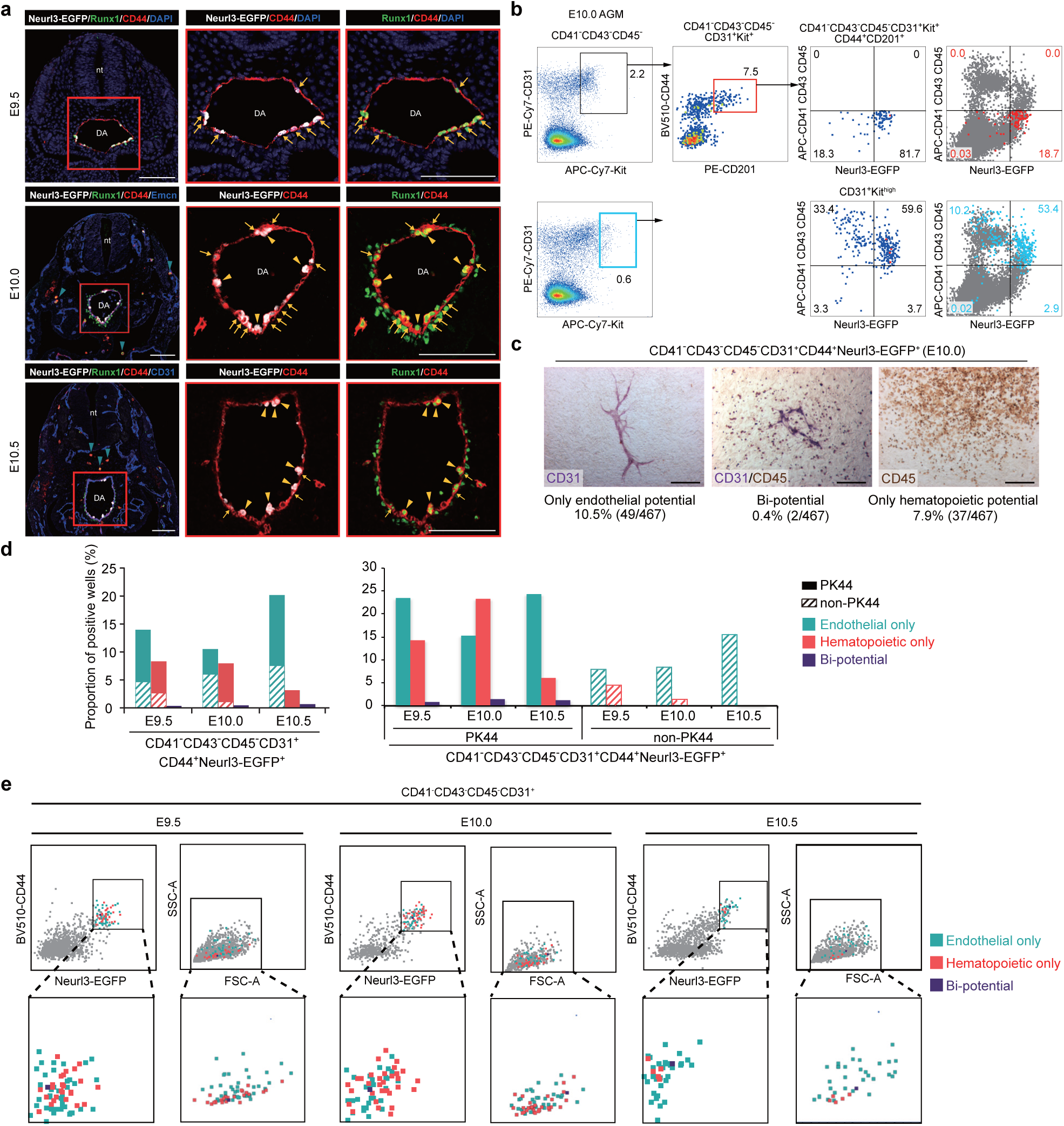
In situ localization and in vitro function of the dynamic HECs marked by Neurl3-EGFP reporter. (a) Representative immunostaining on cross sections at AGM region of E9.5 (upper), E10.0 (middle) and E10.5 (lower) *Neurl3^EGFP/+^* embryos. Arrows indicate Neurl3^+^ aortic ECs. Yellow arrowheads indicate Neurl3^+^ bulging and bulged cells and IAHCs. Aquamarine arrowheads indicate CD44^+^Runx1^+^Neurl3^-^ haematopoietic cells distributed outside the aorta. nt, neural tube; DA, dorsal aorta. Scale bars, 100 μm. (b) Representative FACS analysis of the E10.0 AGM region of *Neurl3^EGFP/+^* embryos. FACS plots to the right showing PK44 cells (red dots, upper) and CD31^+^Kit^high^ cells (blue dots, lower) mapped on, respectively, with their contributions to each gated population indicated. (c) Representative CD31 and CD45 immunostaining on the cultures of single CD41^-^CD43^-^CD45^-^CD31^+^CD44^+^Neurl3-EGFP^+^ cells from E10.0 AGM region of *Neurl3^EGFP/+^* embryos, showing typical morphologies regarding distinct differentiation potentials. Cell frequencies of each kind of potential are also shown. Data are from 5 independent experiments with totally 37 embryos used. Scale bars, 400 μm. (d) Column charts showing the proportions of positive wells in the indicated populations (lower) for each kind of potential. The experiments were performed with CD41^-^CD43^-^CD45^-^CD31^+^CD44^-^Neurl3-EGFP^+^ single cells from E9.5 caudal half or E10.0-E10.5 AGM region of *Neurl3^EGFP/+^* embryos with PK44 indexed. Progenies from PK44 and non-PK44 cells are represented by distinct fill patterns. (e) Expression of CD44 and Neurl3-EGFP and values of FSC-A and SSC-A in the index-sorted single CD41^-^CD43^-^CD45^-^CD31^+^CD44^+^Neurl3-EGFP^+^ cells with differentiation potential based on in vitro functional evaluation. Cells with different kinds of potential are mapped onto the reference FACS plots (grey dots). Solid boxes (left of each stage) indicate the gates of the populations for FACS sorting. The enlarged views of solid boxes are shown below.

Given the lacking of suitable antibodies to directly determine the anatomical distribution of PK44 cells, which have been proven as HSC-competent HECs (Fig. 2c-e), we compared the immunophenotype of PK44 and IAHC cells, known as CD31^+^Kit^high 7^, regarding their relationship with Neurl3-EGFP expression whose localization was clearly defined (Fig. 5a). Both of them were mainly Neurl3-EGFP^+^, with most CD31^+^Kit^high^ cells being CD41/CD43/CD45-positive haematopoietic cells as previously reported ^7^ (Fig. 5b). PK44 showed an expression pattern largely different from CD31^+^Kit^high^ cells, suggestive of their predominant non-IAHC localization (Fig. 5b). The expression of Neurl3-EGFP was completely absent from the sub-aortic mesenchyme, in contrast to the widespread distribution of Runx1 there (Fig. 5a; Supplementary information, Fig. S3e). Although scattered Runx1^+^CD44^+^ round blood cells were easily witnessed, much fewer Neurl3-EGFP-expressed cells outside dorsal aorta were detected, even at E11.0 (Fig. 5a; Supplementary information, Fig. S3e).

As less than half of Neurl3-EGFP^+^ ECs were PK44 cells (Fig. 5b), we next explored the in vitro functional relationship of PK44 and non-PK44 fractions within CD44^+^Neurl3-EGFP^+^ ECs by index-sorting. From E9.5 to E10.5, all three kinds of potential, including endothelial-only, haematopoietic-only, and endothelial-haematopoietic bi-potential, could be detected in CD44^+^Neurl3-EGFP^+^ ECs, with different frequencies (Fig. 5c-e; Supplementary information, Fig. S3f). In E10.5, the potential was remarkably biased to endothelial as compared to E9.5 and E10.0 (Fig. 5d), which should be due to the prompt loss of Neurl3-EGFP-labeled HECs and the possible labeling of some lateAECs by Neurl3-EGFP (Fig. 4a, h). Of note, all three kinds of potential were obviously higher in PK44 than non-PK44 fraction, with endothelial-haematopoietic bi-potential exclusively detected in PK44 cells (Fig. 5d). Therefore, PK44 represented the enriched functional sub-populations within Neurl3-EGFP^+^ HECs. All three kinds of potential did not show an evidently biased distribution regarding CD44 or Neurl3-EGFP expression level by index sorting analysis (Fig. 5e). Interestingly, cells with the haematopoietic rather than endothelial potential intended to have smaller side scatter density on FACS (Fig. 5e).

### Stepwise fate choices of HSC-primed HECs from primitive vascular ECs

In an effort to decipher the stepwise specification of the HSC-primed HECs, we added the immunophenotypic EC samples, from the stage of initial aortic structure formation at E8.0 ^33^ to E9.0, to achieve seamless sampling with continuous developmental stages (Supplementary information, Fig. S4a). All the transcriptomically identified ECs were re-clustered into six clusters, with four of them basically consistent with those previously defined, namely vEC, earlyAEC, lateAEC, and HEC. The newly added samples were mainly distributed into three clusters, vEC, primitive EC (pEC) featured by *Etv2* expression and involving almost all E8.0 cells, and primitive arterial EC (pAEC) given the expression of arterial marker *Gja5* and serving as the earliest arterial EC population, with the latter two clusters as newly identified (Supplementary information, Fig. S4b, c).

Trajectory analysis by Mpath demonstrated two bifurcations along the path from pEC to HEC and revealed a predominant two-step fate choice (Fig. 6a). pEC firstly chose an arterial but not venous fate to become pAEC, then upon maturing into earlyAEC and lateAEC, HEC chose to segregate from the intermediate arterial population earlyAEC (Fig. 6a), in line with the finding that the HEC displayed certain arterial characteristics but was completely devoid of venous feature (Fig. 1f). To decipher the underlying molecular programs for HEC specification, we specifically selected four clusters, excluding vEC and lateAEC branched out from the path from pEC to HEC, and added T1 pre-HSC as the end point for the subsequent analysis (Fig. 6b). Monocle 2 elegantly recapitulated the sequential sampling stages and the deduced cellular evolution upon stepwise hemogenic specification along the inferred pseudotime (Fig. 6c; Supplementary information, Fig. S4d).

**Fig. 6.**
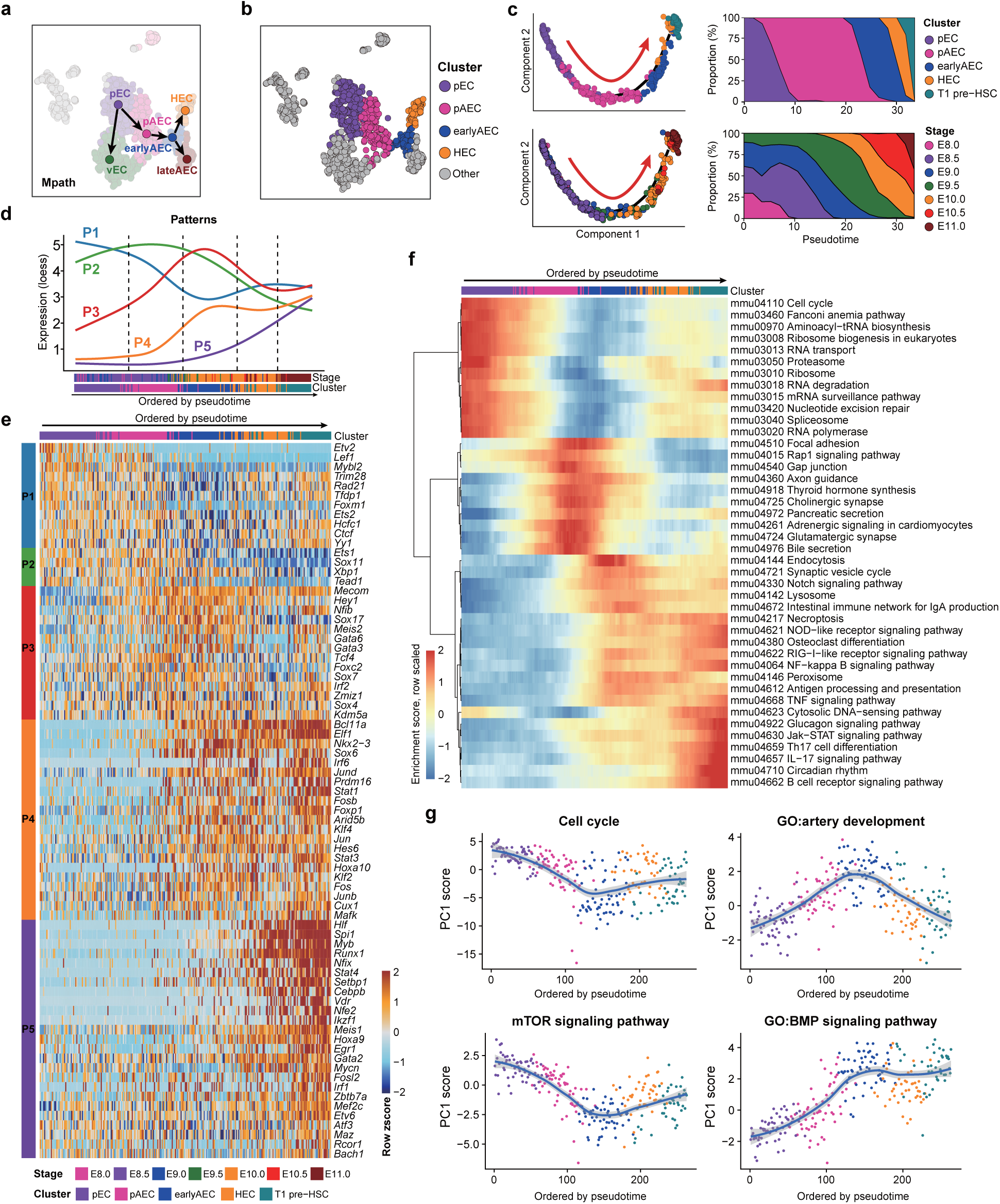
Molecular evolution underlying the specification of HSC-primed HECs from primitive vascular ECs. (a) Trajectory of pEC, vEC, pAEC, earlyAEC, lateAEC and HEC inferred by Mpath. Arrows indicate the development directions predicted by sampling stages. (b) t-SNE plot showing the distribution of the four clusters involved in hemogenic specification. Other cells are in grey. (c) Pseudotemporal ordering of the cells included in the indicated five clusters inferred by monocle 2 (left), with clusters (upper left) and sampling stages (lower left) mapped to it. HEC specification directions are indicated as red arrows. Smooth distribution of clusters (upper right) and sampling stages (lower right) along pseudotime by using Gaussian kernel density estimate are shown to the right. (d) Dynamic changes of five gene expression patterns along the trajectory ordered by pseudotime inferred by monocle 2. For each pattern, principal curves are fitted on expression levels of the genes in that pattern along pseudotemporal order, using local polynomial regression fitting method. Randomly down-sampling is performed in pEC and pAEC clusters for better visualization. (e) Heatmap showing the relative expression of the core TFs which belong to the regulons that the genes within exhibit significant overlapping with the pattern genes. Cells are ordered by pseudotime and TFs are ordered by Patterns. (f) Heatmap showing smoothed (along adjacent 25 cells) and scaled enrichment scores of top 50 KEGG pathways along the order by pseudotime. Pathways are ordered by hierarchical clustering using ward.D method. (g) Scatter plots showing the relative activity levels of pathways or GO terms with loess smoothed fit curves and 95% confidence interval indicated. Relative activity levels are represented by the PC1 scores of expression levels of the genes in a given set. The sign or direction of PC1 is corrected according to positive correlation with averaged expression levels.

We identified totally 2,851 genes whose expression levels were changed significantly among five clusters, which were further grouped into five principal expression patterns along the inferred pseudotime (Fig. 6d; Supplementary information, Fig. S4e and Table S6). In general, genes in Pattern 1 showed the highest expression in pEC, decreased apparently upon arterial specification, whereas slightly increased upon hemogenic specification, which were mainly related to rRNA processing and mitotic nuclear division (Fig. 6d; Supplementary information, Fig. S4e, f). Genes in Pattern 2 showed the highest expression in the initial arterial specification, and the lowest expression upon hemogenic specification, which were mainly related to organization of intra-cellular actin filament and inter-cellular junctions (Fig. 6d; Supplementary information, Fig. S4e, f). Genes in Pattern 3, which were related to endothelium development and cell migration, showed the highest level in earlyAEC, whereas relatively low expression in the upstream pEC and pAEC and downstream HEC and T1 pre-HSC (Fig. 6d; Supplementary information, Fig. S4e, f). Genes in Pattern 4 and Pattern 5 both had the highest expression in the final T1 pre-HSC and both exhibited haematopoiesis-related terms, with those in Pattern 4 reaching the relatively high level from earlyAEC and those in Pattern 5 showing a gradual increase (Fig. 6d; Supplementary information, Fig. S4e,f).

Among the above 2,851 pattern genes, 75 TFs belonged to the core TFs of the regulons where the genes included significantly overlapped with the pattern genes (Fig. 6e; Supplementary information, Table S6). Given the simultaneous co-expression of the core TF and its predicted targets in a given regulon, these core TFs were considered to presumably play a role to drive or orchestrate the dynamic molecular program during HEC specification (Fig. 6e). Most of these TFs belonged to Pattern 4 and Pattern 5, indicating that most activated TFs along HEC specification from primitive vascular ECs were those overrepresented in the final hemogenic and haematopoietic populations (Fig. 6e). We also examined the expression patterns of totally 28 TFs previously reported to have a role in HSPC regeneration in vitro ^34–37^. 19 of these presumed functional TF were dynamically changed and 15 of them were core TFs of regulons, with most of them belonging to Pattern 5 (Supplementary information, Fig. S4g).

We next evaluated the pathway enrichment for each cell to depict dynamic changes at pathway level. The pathways significantly changed among the five candidate clusters showed the dynamic patterns similar to gene expression patterns (Fig. 6f). Among them, cell cycle, ribosome and spliceosome were the pathways that were down-regulated with arterial specification whereas turned to be moderately up-regulated by hemogenic specification (Fig. 6f, g). In contract, several pathways experienced a completely opposite change, such as Rap1 signaling pathway (Fig. 6f). Artery development, together with its pivotal executor Notch signaling pathway ^2, 38^, firstly rose to peak in earlyAEC and then modestly fell down upon hemogenic specification (Fig. 6f, g). Some inflammation related pathways, including NFκB and TNF signaling, were activated from earlyAEC to the final T1 pre-HSC, in line with the notion about the requirement of inflammatory signaling during HSC generation ^39^ (Fig. 6f).

## DISCCUSSION

Here via unbiasedly going through all the relevant EC populations, HSC-primed HECs were transcriptomically identified. More importantly, combining the computational prediction and in vivo functional evaluation, we precisely captured the HSC-competent HECs by a newly constructed fluorescent reporter mouse model, Neurl3-EGFP, and revealed further functionally enriched sub-population within Neurl3-EGFP-labeled ECs by a set of surface marker combination PK44. Serving as the putative marker of HSC-primed HECs ^14, 22^, *Gfi1* was specifically expressed in HEC but not other EC-related populations (Fig. 2h, 4h), supportive of the cluster assignment. Belonging to the gene family of E3 ubiquitin ligases, the expression and role of Neurl3 in spermatogenesis and inflammation has been reported ^40–42^, whereas that relevant to vascular and haematopoietic development remains barely known. Neurl3 was screened out by unsupervised bioinformatics analysis, and fortunately, the expression of which in AGM region was restricted to aorta and largely consistent with that of *Runx1* both transcriptomically (Supplementary information, Fig. S1k) and anatomically (Fig. 5a) regarding endothelial expression. Although highly expressed in tif-HEC, *Runx1* and *Adgrg1* were also highly expressed in the CD45^-^CD43^-^ haematopoietic population (Fig. 4h), which should be the derivatives of non-HSC haematopoiesis. This suggested that they may not distinguish the precursors of HSCs and non-HSCs ^32, 43^, thus *Runx1* and *Adgrg1* were not included in the list of the signature genes of HSC-primed HEC (Fig. 4a). The specificity of *Nerul3* expression related to HSC generation suggests that the Neurl3-EGFP would be a good reporter for the studies of both HSC development and regeneration.

Based on the in vivo functional validation of the HSC-primed HECs and the sampling of continuous developmental stages with intervals of 0.5 days, we had a good opportunity to evaluate the dynamics and functional heterogeneity of these important transient populations. Unexpectedly, the HSC-competent HECs demonstrated a previously unresolved endothelial-haematopoietic bi-potential. The HECs we defined showed a higher enrichment of the expression of key haematopoietic TFs (Supplementary information, Fig. S2d) and of both haematopoietic and endothelial potential than using Runx1 +23GFP^+^ as the maker of HECs ^3^, which might partially explain why the rare endothelial-haematopoietic bi-potential is hardly detected around the timing of HSC emergence in previous report ^3^. Thus, our findings well supplement the functional evaluation of putative HECs, which have a dynamic and transient nature, that without catching the endothelial-haematopoietic bi-potential, it is hard to define a given population to belonging to the ones being experiencing endothelial-to-hemogenic fate determination. Both the constitution and the hemogenic potential of the HSC-competent HECs reached the peak at the time point about 0.5 days before the first HSC emergence, and rapidly decreased thereafter (Fig. 5d; Supplementary information, Fig. S1f and S3f). Interestingly, the endothelial-haematopoietic bi-potential was still maintained until T1 pre-HSC stage at E11.0 (Fig. 3i), when cells have begun to express haematopoietic surface markers (Fig. 3b) and turned the shape into round ^15^. The data suggest that the haematopoietic fate might not have been fixed in T1 pre-HSC, which needs further investigations.

We also precisely decoded the developmental path of HSC-primed HECs from the initially specified vascular ECs, the view of which has been generally neglected previously. We found that the genes and pathways involved in arterial development and Notch signaling were firstly increased and then decreased once upon HEC specification (Fig. 6e, g). Supportively, several seemingly contradictory findings have been reported regarding the role of Notch signaling in HEC specification. For example, activation of arterial program or Notch signaling is known to be required for HEC specification in mouse embryos or generation of HECs with lymphoid potential from human pluripotent stem cells ^44–46^. On the other hand, repression of arterial genes in EC after arterial fate acquisition leads to augmented haematopoietic output ^47^. Noteworthy, we revealed two bifurcates during HSC-primed HEC specification along the path from primitive vascular EC, suggesting two-step fate choice occurred for hemogenic fate settling (Fig. 6a). Serving as the two presumed final fates of earlyAEC, HEC and lateAEC displayed a series of differences (Fig. 1i), which better explains the presumably misinterpreted notion in previous report that arterial ECs and HSCs originate from distinct precursors ^18^. Our findings further emphasize that arterial specification and Notch signaling should be precisely and stepwise controlled for HSC generation. Although both showing obvious similarity regarding the arterial feature and anatomical distribution, the difference between earlyAEC and lateAEC should also be paid attention to as the former but not the latter is the direct origin of the HSC-primed HECs.

It is generally accepted that haematopoietic cells in the IAHCs are proliferative ^48, 49^, within which pre-HSCs are mainly involved ^14, 49^. Supportively, enriched functional T1 pre-HSCs manifested a relatively proliferative status ^11^. On the other hand, slow cycling is witnessed at the base of IAHCs ^49^, and it is suggested that exit from cell cycle is necessary for HEC development and endothelial-to-haematopoietic transition ^44, 50^. Nevertheless, based on the precise recognition of the HSC-primed HECs here, we showed that proliferation was gradually decreased upon arterial specification and maturation, whereas re-activated once the arterial ECs chose a hemogenic fate featured by the simultaneous *Runx1* expression (Fig. 6e-g). The functional requirement of cell cycle control for the specification of the HSC-primed HECs needs to be investigated, which would depend on the initiating cell populations.

We also revealed several similarities regarding the molecular events underlying the development of HSC-primed HECs between in mouse and human embryos we have reported very recently ^19^, including the arterial feature and the overrepresented ribosome and translational activity in the HSC-primed HECs. Such conservation further assures the mouse model as an adequate animal model for HSC development studies. The comprehensive understanding of cellular evolutions and molecular programs underlying the specification of HSC-primed HECs combined with the important spatiotemporal cues *in vivo* will facilitate future investigations directing HSC formation *in vitro* and other related regeneration strategies.

## MATERIALS AND METHODS

No statistical methods were used to predetermine the sample size. The experiments were not randomized at any stage. The investigators were not blinded to allocation during the experiments and outcome assessment.

### Mice

Mice were handled at the Laboratory Animal Center of Academy of Military Medical Sciences in accordance with institutional guidelines. Mouse manipulations were approved by the Animal Care and Use Committee of the Institute. The *Neurl3^EGFP/+^* reporter mouse lines were generated with the CRISPR/Cas9 technique by Beijing Biocytogen. All mice were maintained on C57BL/6 background. Embryos were staged by somite pair (sp) counting: E8.0, 1-7 sp; E8.5, 8-12 sp; E9.0, 13-20 sp; E9.5, 21-30 sp; E10.0, 31-35 sp; E10.5, 36-40 sp; and E11.0, 41-45 sp. In some experiments, caudal half of E10.0 embryo was dissected under heart with limbs removed. AGM region was dissected as reported ^12^. The fluorescent dye Oregon green 488 was purchased from Invitrogen. Staining was performed as previously described ^12^ except that the concentration of staining solution was 5 μmol/L and the time of staining was 3 minutes before washed. Primary embryonic single-cell suspension was acquired by type I collagenase digestion.

### Flow cytometry

Cells were sorted and analyzed by flow cytometers FACS Aria 2 and Calibur (BD Biosciences), and the data were analyzed using FlowJo software (Tree Star). Cells were stained by the following antibodies: B220 (eBioscience, RA3-6B2), CD3 (eBioscience, 145-2C11), CD4 (eBioscience, GK1.5), CD8a (eBioscience, 53-6.7), CD31 (BD or BioLegend, MEC13.3), CD41 (BD or eBioscience, MWReg30), CD43 (BD, S7), CD44 (eBioscience or BioLegend, IM7), CD45.1 (eBioscience, A20), CD45.2 (eBioscience, 104), CD45 (eBioscience, 30-F11), CD144 (eBioscience, eBioBV13), CD201 (eBioscience, eBio1560), Flk1 (eBioscience, Avas12a1), Kit (eBioscience, 2B8), Ly-6G (BioLegend, 1A8), and Mac-1 (eBioscience, M1/70). 7-amino-actinomycin D (7-AAD; eBioscience) was used to exclude dead cells. For index sorting, the FACS Diva 8 “index sorting” function was activated and sorting was performed in single-cell mode.

### OP9-based haematopoietic and endothelial potential assay

Cells were sorted by flow cytometry in single-cell mode and were then plated on the OP9 or OP9-DL1 stromal cells ^51^ in IMDM (Hyclone) containing 15% fetal bovine serum (Hyclone), 1% bovine serum albumin (Sigma), 10 μg/mL insulin (Macgene), 200 μg/mL transferrin (Sigma), and 5.5 x 10^-5^ mol/L 2-mercaptoethanol (Gibco). For the endothelial potential assay, 100 ng/mL rhVEGF-165 (PeproTech) was supplemented. For haematopoietic and endothelial bi-potential assay with 10 cells or single cell plated per well, both 100 ng/mL rhVEGF-165 and 50 ng/mL SCF (PeproTech) were supplemented. After 7 days of co-culture, cells were fixed in 4% paraformaldehyde for 30 minutes and stained with PE-conjugated or purified CD45 antibody (eBioscience, 30-F11 or BD Biosciences) to ascertain the generation of haematopoietic progeny. Subsequently, CD31 (BD Pharmingen, MEC13.3) immunohistochemistry staining was performed using standard procedures, and the formation of CD31-positive tubules in the wells was considered as having endothelial potential.

### OP9-DL1 co-culture and transplantation assay

To investigate the HSC potential of the PK44 population in E10.0 caudal half, male CD45.1/1 and female CD45.2/2 mice were mated to obtain CD45.1/2 embryos. FACS purified cell populations from E10.0 caudal half (CD45.1/2) were plated on the OP9-DL1 stromal cells in α-MEM (Gibco) supplemented with 10% fetal bovine serum (Hyclone) and cytokines (100 ng/mL SCF, 100 ng/mL IL-3 and 100 ng/mL Flt3 ligand, all from PeproTech). After 7 days of co-culture, cells were harvested and then injected into 8-12 weeks female recipients (CD45.2/2) via tail vein, along with 2×10^4^ nucleated fresh bone marrow carrier cells (CD45.2/2) per recipient. Recipients were pre-treated by a split dose of 9 Gy γ-irradiation (^60^Co). Peripheral blood cells of recipients were analyzed by flow cytometry at the indicated time points to determine the chimerism. The recipients demonstrating ≥5% donor-derived chimerism in CD45^+^ cells of peripheral blood were considered as successfully reconstituted. Multi-organ and multi-lineage reconstitution was evaluated as reported ^52^. Totally 1×10^7^ bone marrow cells obtained from the reconstituted primary recipients at 16 weeks post-transplantation were injected into the secondary recipients to investigate HSC self-renewal potential.

To investigate the HSC potential of the CD41^-^CD43^-^CD45^-^CD31^+^CD44^+^Neurl3-EGFP^+^ population in E10.0 caudal half, male *Neurl3^EGFP/+^* reporter mice (CD45.2/2 background) were crossed to female CD45.2/2 mice to generate *Neurl3^EGFP/+^* embryos. Then the co-culture and transplantation strategy were same as mentioned above except that the recipients were female 8-12 weeks CD45.1/2 mice and the carrier cells were obtained from CD45.1/1 mice.

### Immunofluorescence

Embryos were isolated, fixed with 4% paraformaldehyde for 30 minutes to 2 hours at 4°C, embedded in paraffin, and sectioned at 5-6 μ m with Leica RM2235. Sections were deparaffinized with ethanol of gradient concentration, then blocked in blocking solution (Zhongshan golden bridge) for 30 minutes at room temperature, followed by incubation with primary antibodies overnight at 4°C. After 3 washes (3 minutes each) in PBS, sections were incubated with corresponding secondary antibodies (Zhongshan golden bridge) for 30 minutes at room temperature. After 3 washes in PBS, sections were stained with DendronFluor TSA (Histova, NEON 4-color IHC Kit for FFPE, NEFP450, 1:100, 20–60-sec). The primary and secondary antibodies were thoroughly eluted by heating the slides in citrate buffer (pH 6.0) for 10 minutes at 95°C using microwave. In a serial fashion, each antigen was labeled by distinct fluorophores. After all the antibodies were detected sequentially, the slices were finally stained with DAPI. Images were collected by confocal microscope (Nikon Ti-E A1/ ZEISS LSM 880). The primary antibodies were as follows: CD31 (BD Biosciences), CD44 (BD Biosciences), Endomucin (eBioscience), GFP (Cell Signaling), and Runx1 (Abcam).

### Whole-mount Immunofluorescence

The body part between forelimb buds and hindlimb buds of E10.0 embryo was dissected, fixed in 2% PFA/PBS for 20 minutes on ice and dehydrated in graded concentrations of methanol/PBS (50%, 100%; 10 minutes each). To block endogenous peroxidase, samples were bleached in 5% H_2_O_2_ for 1 hour on ice. For staining, the samples were blocked in PBSMT (1% skim milk and 0.4%Triton X-100 in PBS) containing 0.2% BSA for 1 hour at 4℃, incubated with PBSMT containing anti-CD44 (1:25) overnight at 4℃, then washed 3 times in PBSMT each for 1 hour at 4 ℃. The primary antibody was developed by incubating HRP-conjugated anti-rat Ig antibody (1:2000 in PBSMT; Zhongshan golden bridge) overnight at 4℃. After extensive washing with more than 3 exchanges of PBSMT, including the final 20 minutes wash in PBST (0.1% Triton X-100 in PBS) at 4℃, the samples were soaked in DendronFluor TSA (Histova, NEON 4-color IHC Kit for Wholemount/Cytometry, NEWM450) for 10–30 minutes, and hydrogen peroxide was added to 0.03%. The enzymatic reaction was allowed to proceed until the desired color intensity was reached, and the samples were rinsed 3 times in PBST. Finally, the samples were dehydrated in 100% methanol and soaked in graded concentrations of BABB (phenylcarbinol and benzyl benzoate, 1:2)/methanol (50%, 100%; 1 minute each), stored at −20℃ until photographed.

### Single cell RNA-seq library construction

Single cells in good condition were picked into lysis buffer by mouth pipetting. The single cell RNA-seq preparation procedure was based on STRT with some modifications ^53 54 55^. cDNAs were synthesized using sample-specific 25 nt oligo dT primer containing 8 nt barcode (TCAGACGTGTGCTCTTCCGATCT-XXXXXXXX-NNNNNNNN-T25, X representing sample-specific barcode whereas N standing for unique molecular identifiers, UMI, see Table S7) and TSO primer for template switching ^56 57 58^. After reverse transcription and second-strand cDNA synthesis, the cDNAs were amplified by 17 cycles of PCR using ISPCR primer and 3’ Anchor primer (see Table S7). Up to 56 samples were pooled and purified using Agencourt AMPure XP beads (Beckman). 4 cycles of PCR were performed to introduce index sequence (see Table S7). After this step, 400 ng cDNAs were fragmented to around 300 bp by covaris S2. The cDNA was incubated with Dynabeads MyOne^TM^ Streptavidin C1 beads (Thermo Fisher) for 1 hour at room temperature. Libraries were generated using KAPA Hyper Prep Kit (Kapa Biosystems). After adaptor ligation, the libraries were amplified by 7 cycles of PCR using QP2 primer and short universal primer (see Table S7). The libraries were sequenced on Illumina HiSeq 4000 platform in 150bp pair-ended manner (sequenced by Novogene).

### Quantification of gene expression for scRNA-seq data

We used unique molecular identifier (UMI)-based scRNA-seq method to measure the gene expression profiles within individual cells. Raw reads were firstly split by specific barcode attached in Read 2 for individual cells and UMI information was aligned to the corresponding Read 1. Read 1 was trimmed to remove the template switch oligo (TSO) sequence and polyA tail sequence. Subsequently, quality control was conducted to discard reads with adapter contaminants or low-quality bases (N > 10%). Next, the mm10 mouse transcriptome (UCSC) was used to align the clean reads using TopHat (version 2.0.12) ^59^. Uniquely mapped reads were obtained using HTSeq package ^60^ and grouped by the cell-specific barcodes. Transcripts of each gene were deduplicated based on the UMI information, while mitochondrial genes were not included for quantification. Finally, for each gene in each individual cell, the number of the distinct UMIs derived from that gene was regarded as its copy number of transcripts.

### Quality control and normalization of sequencing data

For the 662 sequenced single cells from E9.5-E11.0 embryos of totally 29 embryos, we only retained cells with more than 2,000 genes and 100,000 transcripts detected. Then, 597 cells passed the filter standards. Gene expression levels in each cell were normalized by log_2_(TPM/10+1), where TPM (transcripts-per-million) was calculated as (the number of UMIs of each gene / all UMIs of a given cell) ×1,000,000. Since the UMI number of most of our samples was less than the order of 1,000,000 transcripts, the TPM values were divided by 10 to avoid counting each transcript for several times. On average we detected 7,035 genes (range from 2,266 to 10,843) and 636,418 transcripts (range from 103,793 to 2,959,573) expressed in each individual cell.

Additionally, we also sequenced 96 single cells with a PK44 immunophenotype (CD41^-^CD43^-^CD45^-^CD31^+^CD201^+^Kit^+^CD44^+^) from E10.0 AGM regions of totally 9 embryos, 47 T1 pre-HSCs (CD31^+^CD45^-^CD41^low^Kit^+^CD201^high^) from E11.0 AGM regions of totally 18 embryos, 48 single cells with an immunophenotype of CD41^-^CD43^-^CD45^-^CD31^+^CD44^+^Neurl3-EGFP^+^ from Neurl3-EGFP reporter mouse embryos and 579 single cells from E8.0-E9.0 body regions of totally 24 embryos. The same quality control criteria and normalization method described above were applied to these additional datasets. In total, 1,432 single cells were sequenced and 1,325 cells passed the filter standards and were used for downstream analyses (see Table S1).

### Dimensional reduction and clustering

We used Seurat R package ^61^ (version 2.3.4) for further analyses and exploration of our single cell RNA sequencing data, such as identification of highly variable genes (HVGs) and differentially expressed genes (DEGs), dimension reduction using PCA or t-SNE, unsupervised clustering and so on. A standard analysis process is briefly described below. First, only genes expressed in at least 3 single cells were retained so as to exclude genes that were hardly expressed. Then, FindVariableGenes function was used to select HVGs on log2 (TPM/10+1) transformed expression values. Genes with average expression more than 1 and less than 8 and dispersion greater than 1 were identified as HVGs. To mitigate the effect of cell cycle, HVGs not included in the direct cell cycle GO term (GO:0007049) (Table S7) were used as inputs for PCA dimension reduction. Elbow method was employed to select the top relevant PCs for subsequent t-SNE dimension reduction and graph-based clustering ^28^.

For the initial dataset from E9.5-E11.0 body and DA locations, we select top 15 PCs for clustering using FindClusters with default settings, to obtain 6 major clusters. Negative control cells with a non-EC immunophenotype and cells grouped with these negative control cells were reclassified specifically into the Neg cluster. The remaining cells were assigned as vEC, earlyAEC, lateAEC, HEC and HC clusters based on the clustering results. Next, cells in Neg cluster and cells with *Ptprc* or *Spn* expression level greater than 1 were removed. Then, the filtered initial dataset was used for analyses of subdatasets, including subdividing of HEC cluster, subdividing of eaAEC cluster and in-depth analyses of earlyAEC, lateAEC and HEC clusters. The filtered initial dataset was also included in three combined datasets of combining PK44 cell population, PK44 and T1 pre-HSC cell populations, and PK44 and Neurl3-EGFP cell populations, respectively. Dimension reduction and clustering analyses for subdatasets and combined datasets abovementioned also followed the same procedure as described above. See Table S1 for detailed cell information.

For combined dataset from earlier dataset (E8.0-E9.0 body location) and initial dataset (E9.5-E11.0 body and DA locations), we redid the dimension reduction and clustering analyses over again. Same as the processing of initial dataset, negative control cells with a non-EC immunophenotype and cells grouped with these negative control cells were reclassified manually into Neg cluster. The remaining cells were assigned as vEC, pEC, pAEC, earlyAEC, lateAEC, HEC and HC based on the clustering results. The new clustering results are highly consistent with the previous ones within the common cell populations. Next, cells in Neg cluster and cells with *Ptprc* or *Spn* expression level greater than 1 were removed. Cells in pEC, pAEC, lateAEC and HEC and cells from T1 pre-HSC dataset were retained for further analysis.

### Identification of DEGs

DEGs were identified using FindMarkers or FindAllMarkers functions with default Wilcoxon rank sum test and only genes detected in a minimum fraction of 0.25 cells in either of the two populations were considered. Genes with fold-change ≥ 2 and adjusted *P* value ≤ 0.05 were selected as DEGs.

### Arterial and venous feature score

Arteriovenous marker genes previously known or inferred from the artery development pattern genes, including 10 arterial genes (*Dll4*, *Igfbp3*, *Unc5b*, *Gja4*, *Hey1*, *Mecom*, *Efnb2*, *Epas1*, *Vegfc* and *Cxcr4*) and 3 venous genes (*Nr2f2*, *Nrp2*, and *Aplnr*) ^33, 62–64^, were selected to perform the arteriovenous feature scores. First, we scaled the log_2_(TPM/10+1) expression values of each marker gene to 0-10 scale among all the sample cells after quality control. Second, for each cell, we averaged the scaled values of arterial genes and venous genes, respectively. Third, the averaged values were rescaled to 0-10 scale across all the sample cells to finally achieve the arterial and venous scores. For each population, the arterial and venous scores of all of the cells within the population were average. The 50% confidence ellipses were also calculated to show the main distribution ranges. We chose score value = 5 as the threshold to infer the arterial or venous identity of vascular ECs, as the distribution of individual cells was in line with the notion showing essentially no arterial/venous double positive cells.

### Cell cycle analysis

For cell cycle analysis, cell cycle-related genes consisting of a previously defined core set of 43 G1/S genes and 54 G2/M genes were used ^58, 65^ (see Table S7 for detailed gene lists). We used a way similar to Tirosh, et al. ^66^ to classify the cycling phases of the cells. We calculated the average expression of each gene set as corresponding scores, and manually assigned cells to approximate cell cycle phases based on the scores. Namely, cells with G1/S score < 2 and G2/M score < 2 were assigned as ‘quiescent’, otherwise ‘proliferative’. Among proliferative cells, those with G2/M score > G1/S score were assigned as ‘G2/M’, and those with G1/S score > G2/M score were assigned as ‘G1’ when G2/M score < 2, or as ‘S’ when G2/M score ≥ 2.

### Constructing single cell trajectories

Monocle 2 ^67^ (version 2.6.4) and Mpath ^68^ (version 1.0) were adopted to infer the development trajectory of selected cell populations. Monocle 2 can construct single-cell trajectories and place each cell at its proper position in the trajectory, even a “branched” trajectory corresponding to cellular “decisions”. We followed the official vignette with recommended parameters. Briefly, UMI count data of given cell populations was used as input and genes with more than 1.5 times of fitted dispersion evaluated using dispersionTable function were identified as HVGs. To reduce the influence of cell cycle effect, HVGs not included in the direct cell cycle GO term (GO:0007049) were retained as ordering genes for the subsequent ordering cells.

For Mpath analysis, the log_2_(TPM/10+1) normalized data of HVGs identified by using Seurat method were used as inputs. The cluster labels defined by clustering procedures described above were used as landmark cluster assignment of individual cells. Based on the results of the Mpath analyses, we specified the starting point and developing directions according to the development time and visualized the results on t-SNE plot.

### Patterns of DEGs among multiple clusters

In the case of identification of gene patterns in more than two clusters, analysis of variance followed by Tukey’s HSD test for pairwise comparison was adopted to identify DEGs (genes with adjusted *P* value < 0.05 and fold change > 2 or < 0.5). For identification of patterns in earlyAEC, lateAEC and HEC clusters, only 1,005 DEGs resulted from the pairwise comparisons of earlyAEC and lateAEC and of earlyAEC and HEC were retained. According to the changed directions of HEC and lateAEC as compared to earlyAEC, we could assign these DEGs into 8 patterns as illustrated. Transcription factors network visualization was implemented as follows. First, the average expression values of genes included in each pattern were calculated as their representative expression levels. Then, the representative expression levels of 8 patterns and the expression profile data of transcription factors included in these patterns were combined as input for construction of “signed hybrid” weighted gene co-expression network analysis using WGCNA ^69^. Next, we used 0.01 as adjacency threshold for including edges in the output to export network, which was then imported into Cytoscape^70^ for visualization. We also calculated Pearson correlation coefficient between each transcription factor and the pattern it belongs to.

For identification of patterns in pEC, pAEC, earlyAEC and HEC clusters, all 2,851 DEGs among them were retained. We used ConsensusClusterPlus function with k-means algorithm on top 500 DEGs to achieve five stable clusters. Then all DEGs were reassigned into one of the five patterns according to which pattern had maximum average Pearson correlation coefficient with a given DEG. Note that we used a downsampled dataset in the visualization related to the five patterns in order to show more detailed changes along the development trajectory. Sixty cells were randomly sampled from pEC and pAEC clusters, respectively.

### Identification of HEC signature genes

Firstly, we compared HEC to every cluster to get the overrepresented genes, which were upregulated across each of the other 4 clusters (vEC, earlyAEC, lateAEC and HC) within filtered initial dataset. To make sure the accuracy of HEC overrepresented genes, we used both wilcox and roc method to perform the DEG analysis. Only the DEGs identified by both methods were regarded as HEC overrepresented genes. Finally, 25 cluster HEC overrepresented genes were retained. In order to not only identify the endothelium with hemogenic potential, but also discriminate those HSC-primed hemogenic ECs from yolk sac-derived early haematopoietic populations such as erythro-myeloid progenitors, genes highly expressed in erythro-myeloid progenitors (*Gsta4*, *Spi1*, *Alox5ap* and *Myb*) as reported ^71 72^ were excluded. In addition, genes not highly expressed (log_2_(TMP/10+1) < 2) in HEC or highly expressed (log_2_(TPM/10+1) > 2) in every clusters were also excluded. Finally, eleven HEC overrepresented genes were retained as HEC signature genes.

### SCENIC analysis

SCENIC ^73^ could reconstruct gene regulatory networks from single-cell RNA-seq data based on co-expression and DNA motif analysis. Here, we used SCENIC R package (version 1.1.1-9) to identify refined regulons, each of which represented a regulatory network that connects a core TF with its target genes. We followed the “Running SCENIC” vignette in the R package with default settings. We identified 507 unique regulons, among which 75 regulons significantly overlapped with the 2851 significantly changed genes were retained. Fisher’s exact test was employed for estimate the statistical significance of their overlaps. The 75 core TFs were considered as putative driving force to orchestrate the dynamic molecular program during HEC specification, given the simultaneous co-expression of the core TF and its predicted targets in a given regulon.

### Gene set variation analysis

Through gene set variation analysis, gene-level expression profiles could be transformed into pathway-level enrichment score profiles using GSVA R package ^74^ coupled with KEGG pathways ^75^. We used ssgsea method ^76^ to estimate gene-set enrichment scores of each cell. Two-sample Wilcoxon test was employed to find differentially enriched pathways between involved clusters. Adjusted *P* value < 0.05 was considered statistically significant.

### TFs and cell surface molecules

Genes were marked as TFs according to 1,485 TFs included in AnimalTFDB 2.0 ^77^, and marked as surface molecules according to 871 high-confidence surfaceome proteins identified in Cell Surface Protein Atlas ^78^. See Table S7 for the detailed gene lists.

### Statistical analysis

All statistical analyses were conducted in R version 3.4.3. Two-sample Wilcoxon Rank Sum test was employed for comparisons of gene numbers, transcript counts, or gene expression levels between two clusters of cells. We referred to statistically significant as *P* < 0.05 (if not specified). Network enrichment analyses and gene ontology biological process enrichment analyses were performed using Metascape ^79^ (http://metascape.org) and clusterProfiler ^80^, respectively.

### Data and Code Availability

The scRNA-seq data has been deposited in the NCBI Gene Expression Omnibus, the accession number for the data is pending. Code is available on reasonable request.

---

**Fig. S1.**
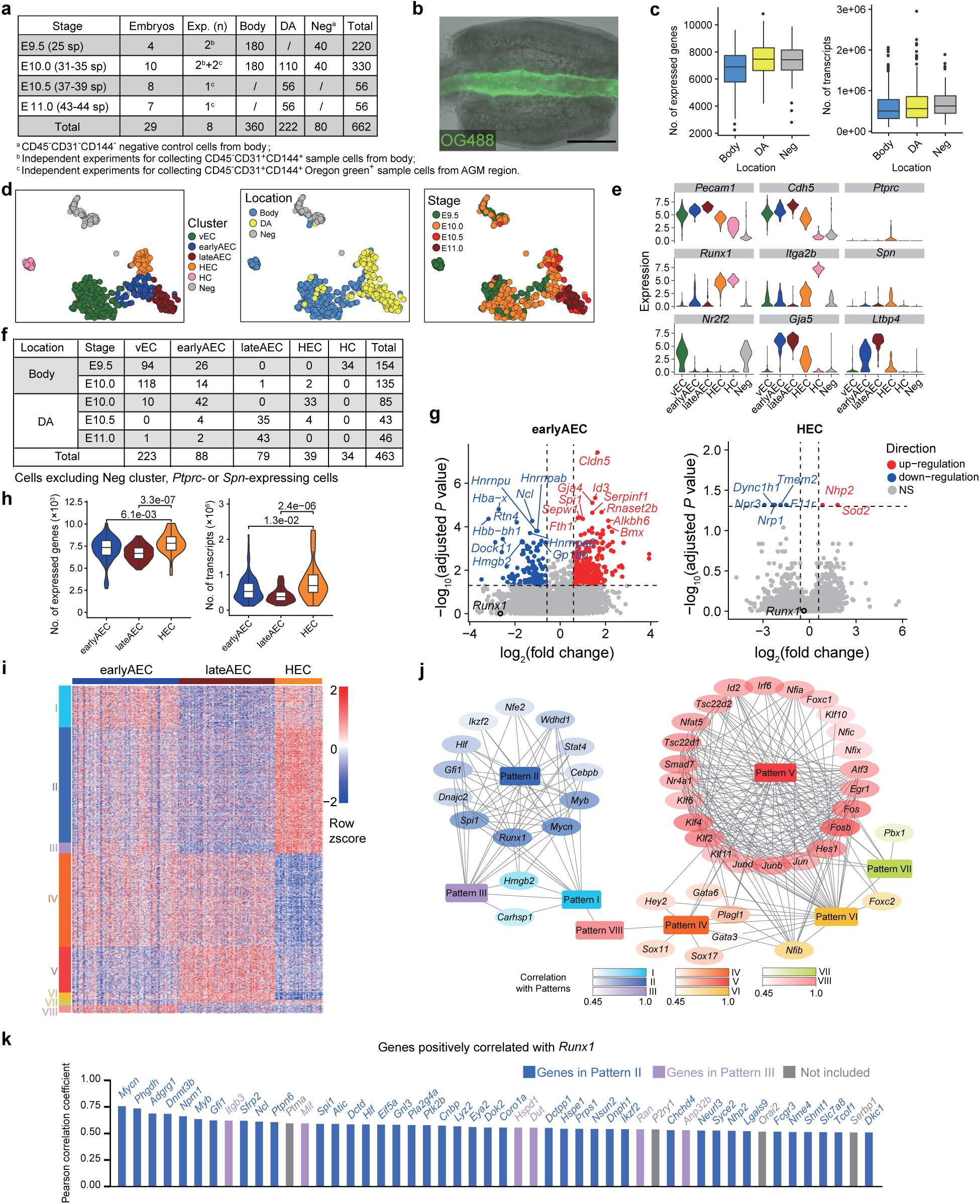
Information, clustering of initial dataset and molecular characteristics of major clusters. (a) Embryo, independent experiment, and cell number information for scRNA-seq. DA, dorsal aortic luminal layer of AGM region. (b) Whole-mount image of the E11.0 AGM region labeled with Oregon Green 488. (c) Boxplots showing the number of genes (left) and transcripts (right) in each single cell of different locations. (d) t-SNE plots with clusters (left), sampling locations (right) and embryonic stages (right) mapped onto it. (e) Violin plots showing the expression levels of indicated genes in six clusters identified in the initial dataset. (f) Cell number information of the spatiotemporal distribution of distinct clusters. (g) Volcano plots showing differentially expressed genes (marked as blue or red) between two sub-clusters by forced clustering in earlyAEC and HEC, respectively. Top 10 (earlyAEC) or all (HEC) differentially expressed genes are indicated. *Runx1* and *Kit* are also indicated. (h) Violin plots showing the number of genes (left) and transcripts (right) in each single cell of the indicated clusters. Wilcoxon Rank Sum test is employed to test the significance of difference and *P* values are indicated for the comparison. *P* < 0.05 is considered statistically significant. (i) Heatmap showing the relative expression levels of genes in eight patterns among earlyAEC, lateAEC and HEC. (j) Network view of TFs positively correlated with the gene expression patterns. A deeper background color of the gene name indicates a higher positive correlation of the TF to that expression pattern. (k) Bar chart showing the top 50 genes positively correlated with Runx1 within cell population including earlyAEC, lateAEC and HEC. Genes included in the patterns identified above are marked as indicated.

**Fig. S2.**
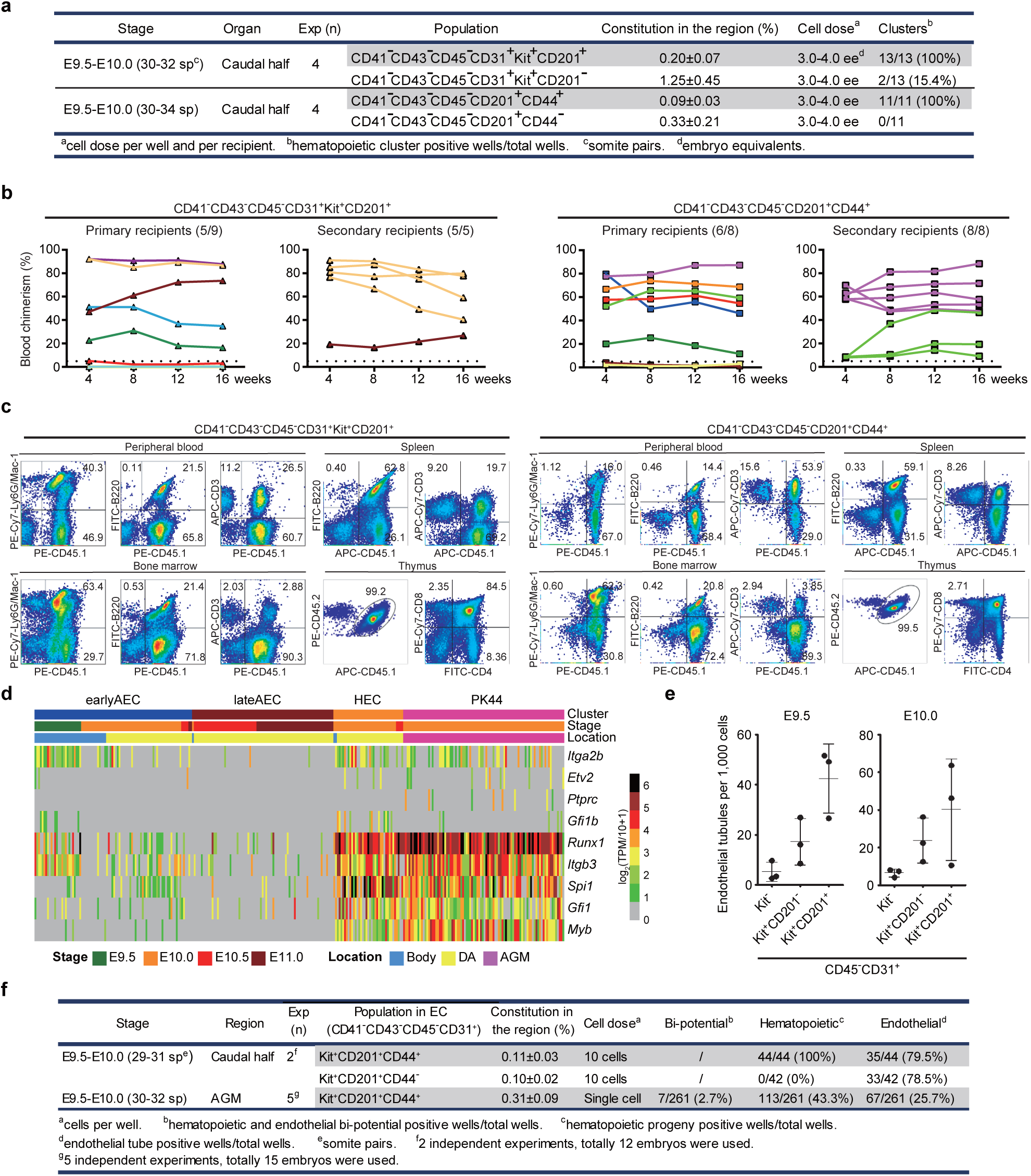
Identification of the HSC-competent and endothelial-haematopoietic bi-potent HECs. (a) Detailed information of the co-culture/transplantation assays performed with E9.5-E10.0 caudal half cells. (b) Blood chimerism of the primary and secondary recipients at 4-16 weeks post-transplantation. The primary recipients were transplanted with the derivatives of the indicated cell populations from the caudal half of E9.5-E10.0 embryos. The paired primary and corresponding secondary repopulated mice are show as the same symbol and color. Numbers of repopulated/total recipients are shown in the brackets. Only the recipients survived to 16 weeks post-transplantation are shown. (c) FACS plots showing representative primary recipients with long-term (16 weeks), multi-organ and multi-lineage repopulations transplanted with the derivatives of the indicated cell populations from the caudal half of E9.5-E10.0 embryos. Donor-derived (CD45.1+CD45.2+) myeloid (Gr-1+/Mac-1+), B lymphoid (B220+), and T lymphoid (CD3+) cells in multiple haematopoietic organs are shown. (d) Heatmap showing the expression of selected genes in earlyAEC, lateAEC, HEC and PK44 populations. Note the similarity of expression patterns between HEC and PK44. (e) Graph showing the endothelial potential of different cell populations in E9.5-E10.0 body part of embryo proper. Cells with indicated immunophenotype were isolated by FACS, co-cultured with OP9 stromal cells for 7 days, and stained with CD31 to identify the endothelial tubes. Data are means ± s.d.. For E9.5 embryos, data are from 3 independent experiments with 6-9 embryo equivalents pooled for each experiment. For E10.0 embryos, data are from 3 independent experiments with 8-9 embryo equivalents pooled for each experiment. (f) Detailed information of endothelial-haematopoietic bi-potential induction assays performed with cells from E9.5-E10.0 caudal half or AGM region.

**Fig. S3.**
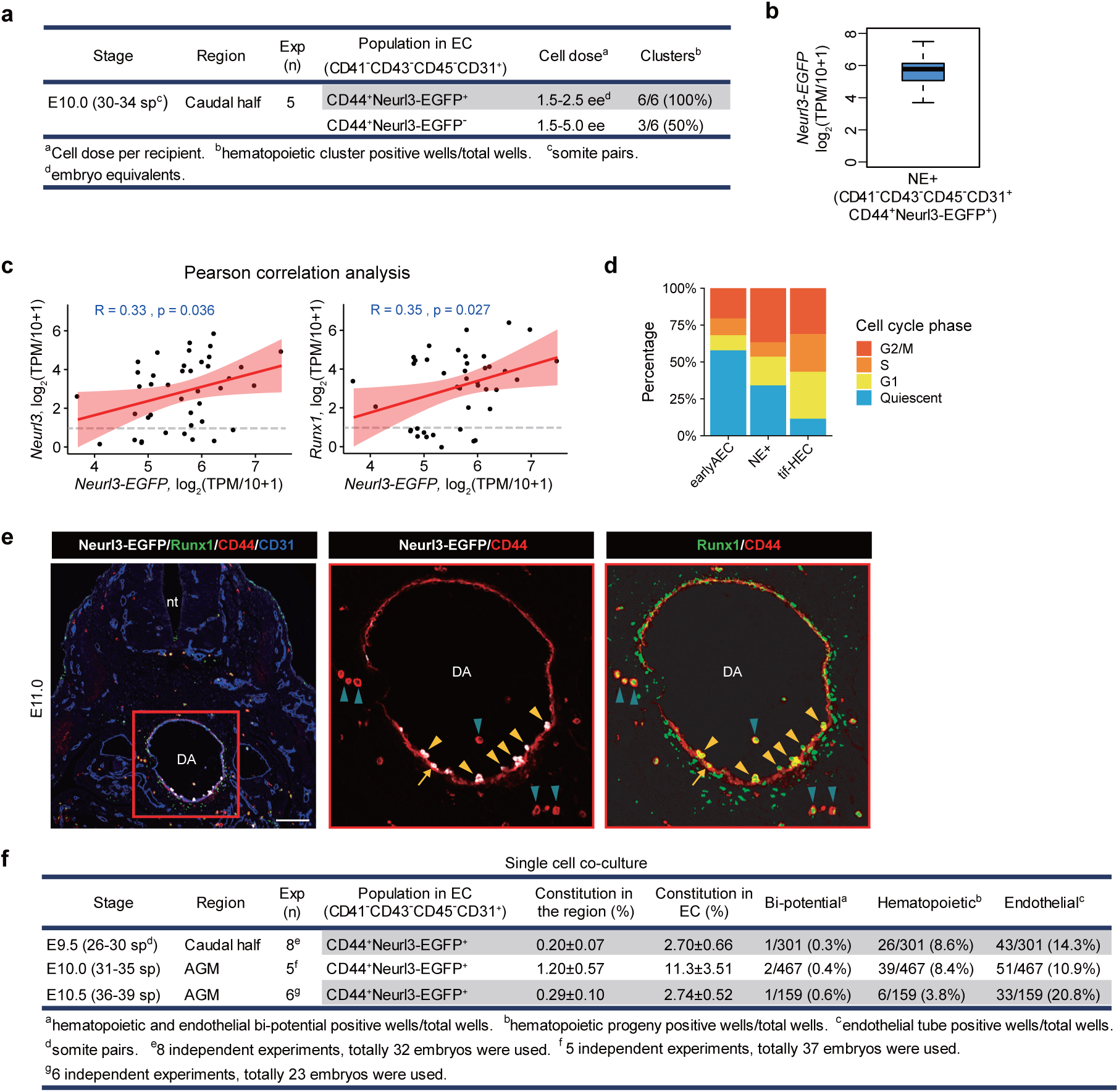
Identification of the HSC-competent HECs marked by Neurl3-EGFP reporter. (a) Detailed information of the co-culture/transplantation assays performed with the caudal half cells from E10.0 *Neurl3^EGFP/+^* embryos. (b) Boxplot showing the transcriptional expression level of *EGFP* in NE+ cell population, NE+, CD41^-^CD43^-^CD45^-^CD31^+^CD44^+^Neurl3-EGFP^+^ population from AGM region of E10.0 *Neurl3^EGFP/+^* embryos. (c) Scatter plots showing correlation of the expression of *EGFP* with that of *Neurl3* and *Runx1*, respectively. Fitted line and 95% confidence interval are shown in red. Pearson correlation coefficients and *P* values are also shown in blue text. (d) Stacked bar chart showing the constitution of different cell cycle phases in the indicated clusters. (e) Representative immunostaining on cross sections at AGM region of E11.0 *Neurl3^EGFP/+^* embryos. Arrow indicates Neurl3^+^ aortic ECs; Yellow arrowheads indicate Neurl3^+^ bulging and bulged cells and IAHCs. Aquamarine arrowheads indicate CD44^+^Runx1^+^Neurl3^-^ haematopoietic cells distributed outside the aorta. nt, neural tube; DA, dorsal aorta. Scale bars, 100 μm. (f) Detailed information of endothelial-haematopoietic bi-potential induction assays performed with cells from E9.5 caudal half and E10.0-E10.5 AGM region of *Neurl3^EGFP/+^* embryos.

**Fig. S4.**
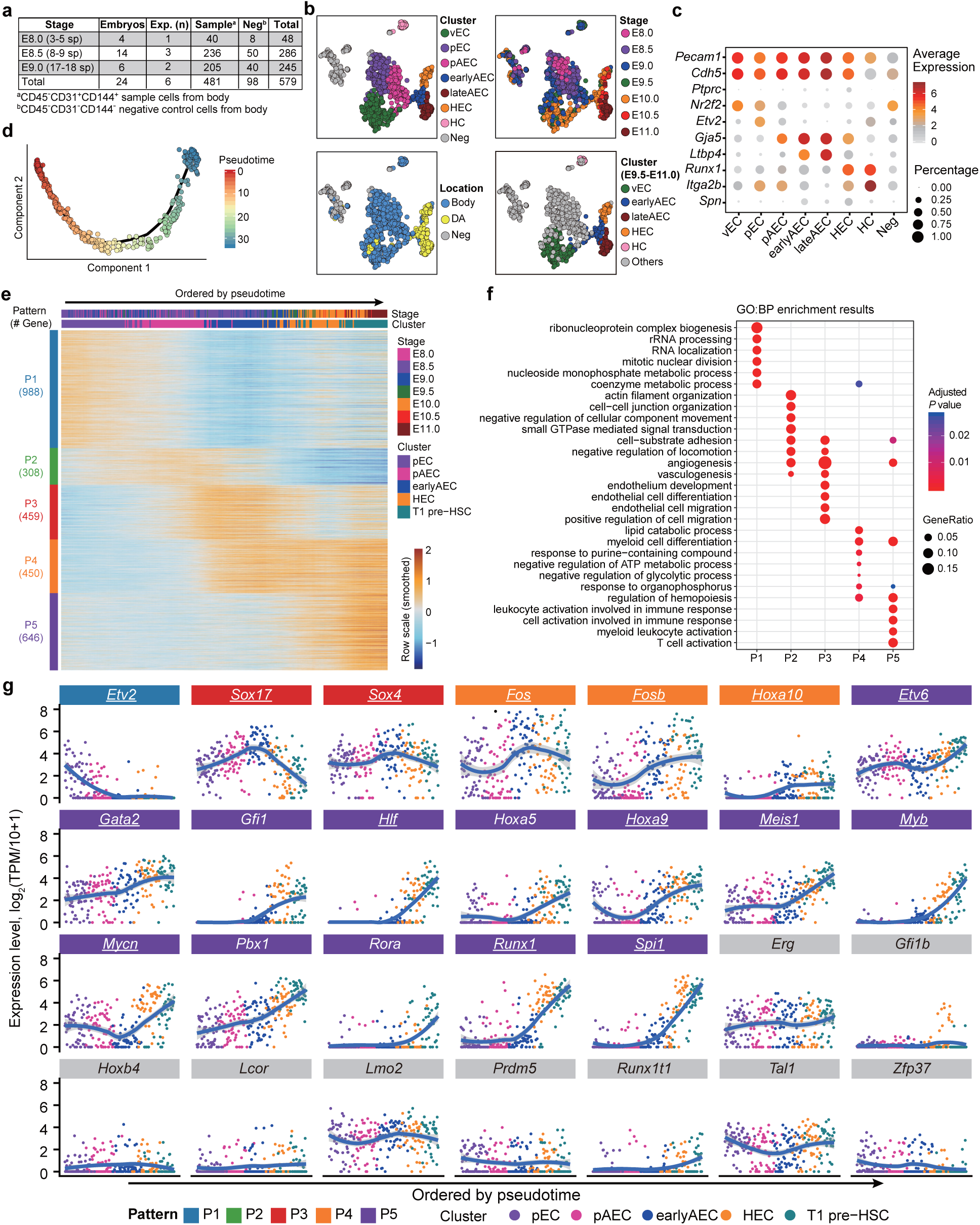
Molecular programs from primitive vascular ECs to HSC-primed HECs. (a) Embryo, independent experiment, and cell number information for additional scRNA-seq. sp, somite pairs. (b) t-SNE plots with clusters (upper left), sampling locations (lower left), embryonic stages (upper right) and clusters previously defined (lower right) mapped onto it. (c) Dot plot showing the average and percentage expression of selected marker genes in the indicated clusters. (d) Pseudotemporal ordering of the cells involved in HEC specification, including those in pEC, pAEC, earlyAEC, HEC, and T1 pre-HSC, inferred by monocle 2, with pseudotime mapped to it. (e) Heatmap showing smoothed and scaled expression levels of 2,851 pattern genes. Genes are ordered by patterns. Cells are ordered by pseudotime. (f) Dot plot showing the top six enriched Gene Ontology biological process (GO:BP) terms for each pattern. Dot color indicates statistical significance of the enrichment and dot size represents the fraction of genes annotated to each term. (g) Scatter plots showing the expression levels of the TF genes previously reported to be functional in HSPC regeneration along the pseudotemporal order with loess smoothed fit curves and 95% confidence interval indicated. The patterns to which the genes belong are indicated by different fill colors. The core TFs of the significantly overlapped regulons are underlined.

